# Tempo of degeneration across independently evolved non-recombining regions

**DOI:** 10.1101/2021.07.20.453045

**Authors:** Fantin Carpentier, Ricardo C. Rodríguez de la Vega, Paul Jay, Marine Duhamel, Jacqui A. Shykoff, Michael H. Perlin, R. Margaret Wallen, Michael E. Hood, Tatiana Giraud

**Author notes:** These authors co-supervised the study (Michael E. Hood, Tatiana Giraud). **Corresponding authors:** Tatiana Giraud and Ricardo C. Rodríguez de la Vega Laboratoire Ecologie, Systématique et Evolution, Bâtiment 360, Université Paris-Saclay, 91400 Orsay France.

## Abstract

Recombination is beneficial over the long term, allowing more effective selection. Despite long-term advantages of recombination, local recombination suppression can evolve and lead to genomic degeneration, in particular on sex chromosomes. Here, we investigated the tempo of degeneration in non-recombining regions, i.e., the function curve for the accumulation of deleterious mutations over time, leveraging on 22 independent events of recombination suppression identified on mating-type chromosomes of anther-smut fungi, including newly identified ones. Using previously available and newly generated high-quality genome assemblies of alternative mating types of 13 *Microbotryum* species, we estimated degeneration levels in terms of accumulation of non-optimal codons and non-synonymous substitutions in non-recombining regions. We found a reduced frequency of optimal codons in the non-recombining regions compared to autosomes, that was not due to less frequent GC-biased gene conversion or lower ancestral expression levels compared to recombining regions. The frequency of optimal codons rapidly decreased following recombination suppression and reached an asymptote after ca 3 Mya. The strength of purifying selection remained virtually constant at d_N_/d_S_ = 0.55, *i.e*. at an intermediate level between purifying selection and neutral evolution. Accordingly, non-synonymous differences between mating-type chromosomes increased linearly with stratum age, at a rate of 0.015 per MY. We thus develop a method for disentangling effects of reduced selection efficacy from GC-biased gene conversion in the evolution of codon usage and we quantify the tempo of degeneration in non-recombining regions, which is important for our knowledge on genomic evolution and on the maintenance of regions without recombination.

## Introduction

Recombination is beneficial over the long term in most situations, allowing higher selection efficacy and therefore more rapid adaptation. Meiotic recombination, through crossing-over resulting in the reciprocal exchange of DNA segments between two homologous chromosomes, shuffles allelic combinations, such that more beneficial combinations might be formed (Fisher 1930; Müller 1932) and selection at different loci can be decoupled (Bernstein, et al. 1981; Otto and Lenormand 2002). Over long time scales, crossover events indeed occur along chromosomes and selection can apply at each locus independently. Recombination prevents selection interference between loci (Hill and Robertson 1966), thereby facilitating the purge of deleterious mutations. In the absence of recombination, selection on strongly beneficial alleles can increase the frequency of linked deleterious alleles through genetic hitchhiking (Maynard Smith and Haigh 1974). Recombination also prevents the accumulation of deleterious mutations by generating chromosomes with fewer deleterious mutations than parental ones, thus avoiding the increase in the frequency of genotypes with a higher number of harmful mutations, known as Muller’s ratchet (Fisher 1930; Muller 1932).

Despite these long-term advantages of recombination, local recombination suppression can be selected for, thereby maintaining beneficial allelic combinations and leading to a genetic structure known as a supergene with multiple linked genes that are transmitted as a single locus (Gutiérrez-Valencia, et al. 2021; Schwander, et al. 2014). Examples of complex phenotypes controlled by supergenes include wing patterns in *Heliconius* butterflies (Nadeau 2016; Saenko, et al. 2019), social systems in ants (Wang, et al. 2013) and mating compatibility systems, such as sex chromosomes or mating-type loci (Charlesworth and Charlesworth 1978; Hartmann, et al. 2021; Ohno 1967; Westergaard 1958). The suppression of recombination can thus arise and be maintained by selection, but the corresponding genomic regions then accumulate deleterious mutations through interferences between loci and Muller's ratchet (Bachtrog 2013; Charlesworth and Charlesworth 2000; Gutiérrez-Valencia, et al. 2021; Jay, et al. 2021a; Stolle, et al. 2019). Typical deleterious mutations include non-synonymous substitutions (Berlin and Ellegren 2006; Brion, et al. 2020; Hough, et al. 2014; Nicolas, et al. 2004; Papadopulos, et al. 2015; Whittle, et al. 2011) which alter the amino-acid sequence of a protein and may result in protein dysfunction. Deleterious mutations also include frameshift mutations, gene losses and substitutions leading to suboptimal gene expression level (Bachtrog 2005; Bartolomé and Charlesworth 2006; Steinemann and Steinemann 1992). Such deleterious mutations in sex chromosomes may be responsible for genetic diseases (Bianchi 2009; Lee, et al. 2016). Genomic rearrangements (Badouin, et al. 2015),epigenetic modifications (Yan, et al. 2005) and transposable elements (Steinemann and Steinemann 1992; Stolle, et al. 2019) also accumulate in non-recombining regions; they are not *per se* deleterious mutations but may disrupt gene function or expression and impose strong energetic costs (Hollister and Gaut 2009; Li, et al. 2016).

A less studied type of deleterious mutation concerns some synonymous substitutions. Synonymous substitutions are usually considered neutral because they do not alter the amino-acid sequences of proteins. However, synonymous codons are often not used randomly (Duret 2002; Sharp, et al. 1995), some codons being preferentially used over their synonymous alternative codons. This preference is thought to result from selection on either the rate or the accuracy of translation (Machado, et al. 2020), with the preferred codons called optimal codons. Optimal codons tend to correspond to the most abundant tRNAs (Ikemura 1981; Post, et al. 1979),and are most frequent in highly expressed genes for which it is most important to avoid limitations in tRNAs (Novoa and Ribas de Pouplana 2012; Zhou, et al. 2016; Wint et al. 2022).

The existence of preferred codons among synonymous alternatives leads to a codon usage bias, which is directly proportional to the recombination rate along genomes (Kliman and Hey 1993). Such deviations from preferred codon usage is thought to result from low selection efficacy in regions with low or no recombination (Bartolomé and Charlesworth 2006). Compared to recombining regions, lower frequencies of optimal codons have been reported in the non-recombining region of the mating-type chromosomes in the fungus *Neurospora tetrasperma* (Whittle, et al. 2011), in the non-recombining dot chromosomes in *Drosophila americana* (Betancourt and Welch 2009) and in twelve other *Drosophila* species (Vicario, et al. 2007).However, codon biases could also result from the biased mutational pattern caused by GC-biased gene conversion. Occurring in regions of frequent recombination, gene conversion results from the resolution of base-pair mismatches in the heteroduplex of the recombination process (Duret and Galtier 2009). GC-biased gene conversion occurs in bacteria (Lassalle, et al. 2015) and eukaryotes (Duret and Galtier 2009; Marais 2003; Pessia, et al. 2012; Weber, et al. 2014). Thus, reduced codon bias in non-recombining regions would be consistent with either fewer GC-biased gene conversion events or less efficient selection, and the respective effects of these two processes may be difficult to disentangle. Few studies have tried to distinguish the roles of relaxed selection versus GC-biased gene conversion (Kostka, et al. 2012) and no study to our knowledge specifically attempted to control for the effect of GC-biased gene conversion in assessing the effect of the lack of recombination on the frequency of optimal codons.

While the accumulation of deleterious mutations in non-recombining regions has been reported in numerous systems, only a few studies with limited data have addressed the question of the tempo of degeneration *i.e*. the rate at which deleterious mutations accumulate over time (Charlesworth 2021). Theoretical works based on the mammalian Y-chromosome predict that degeneration through gene losses should occur rapidly in young non-recombining regions, mainly due to Muller’s ratchet and background selection, and more slowly in older non-recombining regions in which genetic hitchhiking is a major driving force (Bachtrog 2008, 2013).Such patterns of slower gene loss rate with increasing age of recombination suppression have been observed not only in mammals (Bellott, et al. 2014; Hughes and Rozen 2012) but also in the plant *Silene latifolia*, where the sex chromosomes underwent two successive recombination suppression events, *i.e*., evolutionary strata (Lahn and Page 1999). These have been losing genes at estimated rates of 4.64% and 6.92% per million year (MY) for the older and younger evolutionary strata, respectively (Krasovec, et al. 2018). However, a formal study of the tempo of degeneration requires data from a large number of independent events of recombination suppression of varying ages, which has not been available so far (Charlesworth 2021).Recombination suppression has however evolved numerous times in a wide range of organisms, even sometimes multiple independent times within the same genus (Ma and Veltsos 2021; Mrackova, et al. 2008). Moreover, the evolutionary strata present in some sex chromosomes represent stepwise extension events of recombination suppression that occurred at different time points; *e.g*., the four successive steps of recombination suppression on the mammalian Y chromosome (Lahn and Page 1999; Ross, et al. 2009; Skaletsky, et al. 2003).

In the fungal genus *Microbotryum*, recombination suppression has evolved independently in several species, leading to mating-type chromosomes of different ages, some showing stepwise extension of recombination suppression and thereby the formation of evolutionary strata (Branco, et al. 2017; Branco, et al. 2018; Carpentier, et al. 2019; Duhamel, et al. 2022). In these pathogenic anther-smut fungi, mating occurs mostly between haploid cells resulting from the same meiosis event (*i.e*., intra-tetrad mating or automixis (Hood and Antonovics 2000)) and between cells bearing distinct alleles at the two mating-type loci, *i.e*. the PR locus (carrying pheromone and pheromone receptor genes controlling gamete fusion) and the HD locus (carrying homeodomain genes whose proteins undergo dimerization to induce post-mating growth). Recombination suppression linking a few genes within each of the two mating-type loci is required for ensuring correct mating-type functions, and is the ancestral condition in the basidiomycete fungi (Coelho, et al. 2017; Hartmann, et al. 2021). In *Microbotryum* and some other selfing basidiomycetes, the linkage of the two mating-type loci to each other or the linkage of each mating-type locus to its respective centromere has been favoured by selection (Branco, et al. 2017; Branco, et al. 2018; Carpentier, et al. 2019; Hartmann, et al. 2021).Indeed, under intra-tetrad mating these linkage relationships increase the odds of gamete compatibility (producing only two inter-compatible mating-type phenotypes among haploid meiotic products) compared to independent segregation (producing up to four different mating types among gametes following a meiosis). Across the *Microbotryum* genus, five independent recombination suppression events have been shown to link the PR and HD loci to each other (Branco, et al. 2017; Branco, et al. 2018). The two mating-type loci were ancestrally located on different chromosomes, and multiple PR-HD linkage events across lineages involved distinct rearrangements (fusion and/or fission) of the ancestral PR and HD chromosomes (Branco, et al. 2018). Two additional recombination suppression events in other *Microbotryum* species independently linked each of the PR and HD loci to its respective centromere (Carpentier, et al. 2019). Furthermore, extensions of recombination suppression have occurred stepwise and independently in several *Microbotryum* species, forming evolutionary strata of different ages (Branco, et al. 2017; Branco, et al. 2018). The evolutionary strata linking HD and PR loci to each other or to centromeres were all called “black evolutionary strata”, though they corresponded to several independent events, often trapping different sets of genes. In some cases, the full recombination cessation linking the HD and PR loci occurred in several steps and the corresponding strata were distinguished as grey and black (Duhamel, et al. 2022). Extensions of recombination suppression to non-mating-type genes were called coloured evolutionary strata (Branco, et al. 2017; Branco, et al. 2018). The oldest evolutionary strata extending recombination suppression to non-mating-type genes occurred around each of the PR and HD loci; these strata likely evolved before the radiation of the *Microbotryum* genus because they are shared by all species studied so far. Other species-specific strata arose more recently and distal to the fused PR-HD loci. All species retain a recombining region at least at one extremity of their mating-type chromosome, called the pseudo-autosomal region (PAR).

The non-recombining regions in the *Microbotryum* mating-type chromosomes seem to have accumulated multiple footprints of degeneration, in the form of higher rate of non-synonymous substitutions, transposable element accumulation, gene losses and reduced gene expression (Fontanillas, et al. 2015). However, our understanding of the mating-type chromosome content and evolution has changed since, with improved assembly quality and the discovery of the independent origins of the non-recombining regions across species as well as multiple evolutionary strata within species. High-quality assemblies permitted us to detect independent recombination suppression in five *Microbotryum* species, with accompanying degeneration including gene gain/loss and transposable element accumulation (Branco, et al. 2018). Regions with older recombination suppression appeared more degenerated (Branco, et al. 2018), but the tempo of degeneration was not studied. Degeneration in terms of gene expression level was also found based on the high-quality assembly of *M. lychnidis-dioicae*, with reduced expression being associated with different types of degenerative mutations (Ma, et al. 2020).

Here, we investigated the tempo of degeneration in non-recombining regions, *i.e*., the relationship between deleterious mutation accumulation and time since recombination cessation, taking advantage of the multiple independent linkage events and of evolutionary strata of different ages in the anther-smut fungi. We first identified three additional independent events of recombination suppression that arose from linking the PR and HD loci on the mating-type chromosomes, using the genome assemblies reported here for the first time of *M. v. tatarinowii, M. v. melanantha* and *M. coronariae*. We also identified two additional old strata in the large non-recombining mating-type chromosome of *M. v. paradoxa*. Using high-quality genome assemblies of alternative mating types of 13 *Microbotryum* species we estimated the degeneration levels in terms of the accumulation of non-optimal codons and non-synonymous substitutions in 22 independent evolutionary strata of different ages. We identified optimal codons as those enriched in highly expressed autosomal genes and tested whether these were less frequent in non-recombining regions after controlling for the following possible confounding effects: i) lower GC-biased gene conversion in non-recombining regions, by comparing GC content in coding versus non-coding regions, because selection on codon usage acts only in exons while GC-biased gene conversion also impacts introns and inter-genic sequences, and ii) lower ancestral expression levels in non-recombining regions, taking as a proxy the expression level in a closely related outgroup without recombination suppression. We then investigated the relationship between degeneration level (in terms of non-synonymous substitution accumulation and decrease in frequency of optimal codons) and the time since recombination suppression in the different evolutionary strata and species. We found that the frequency of optimal codons rapidly decreased following recombination suppression and reached an asymptote at ca 3 MY. The strength of purifying selection remained virtually constant at an intermediate level between purifying selection and neutral evolution: the ratio of non-synonymous over synonymous differences between alleles on alternative mating-type chromosomes (d_N_/d_S_) indeed remained at 0.55. Accordingly, the non-synonymous mutation differences (d_N_) between mating-type chromosomes increased linearly with stratum age. Understanding the tempo of degeneration is important for our knowledge on genomic evolution, in particular for the maintenance of regions without recombination, that can be associated with lower fitness and human genetic diseases (Wilson 2021).

## Results

### PR-HD linkage in three new *Microbotryum* genomes

We sequenced (long-read Pacific Bioscience technology) and assembled the genomes from haploid cells bearing opposite mating types (called a_1_ and a_2_) of a single diploid individual, for the following species: *Microbotryum violaceum* (*s.l*.) parasitizing *Silene melanantha* (called *M. v. melanantha*), *M. violaceum* (*s.l*.) parasitizing *S. tatarinowii* (called *M. v. tatarinowii*) and *M. coronariae* parasitizing *Silene flos-cuculi* (Syn. *Lychnis flos-cuculi*) (Figure 1A; Table S1). In most species, the mating-type chromosomes were assembled in one or few contigs, in the latter case being broken not at the same position in the two mating-type chromosomes within diploid individuals (Figure S1), which allowed joining contigs. We reconstructed the evolutionary history of mating-type chromosomes by comparing their genome structure to those of *M. intermedium*, taken as a proxy for the genomic ancestral state, following previous studies (Branco, et al. 2017; Branco, et al. 2018). In *M. intermedium*, the PR and the HD mating-type loci are located on distinct mating-type chromosomes (Figure S2). In the case of *M. coronariae*, we could not assess the respective positions of several contigs within the non-recombining region (Figure S1), but this did not affect the general scenario of rearrangements (Figure 1A) as these contigs could be assigned to the non-recombining region given the rearrangements between mating types and could be assigned to ancestral mating-type chromosome arms (Figure S1). For reconstructing the scenarios of chromosomal rearrangements, mating-type chromosome fissions (always at centromeres) were inferred by assessing what chromosome arms became autosomes (*i.e*., completely collinear between mating types and assembled separately from the mating-type chromosomes in both haploid genomes; Figure S1). The orientations of ancestral mating-type chromosome fusions were inferred by assessing on Figure S1 what edges of ancestral mating-type chromosomes remained recombining (*i.e*. became PARs, the other edge or centromere then corresponding to the fusion point).

**Figure 1:**
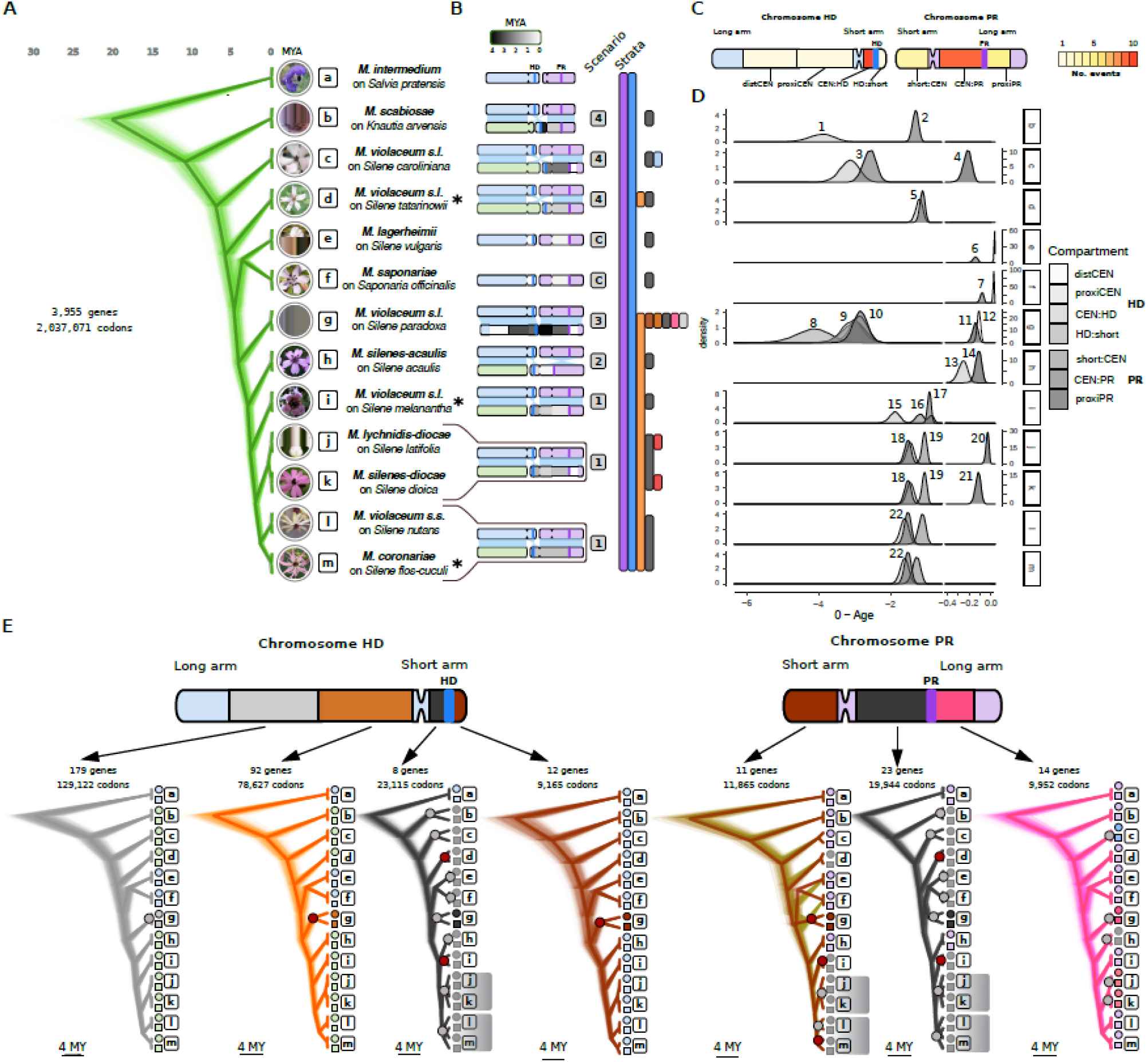
Timetrees and rearrangement scenarios of recombination suppression events in the *Microbotryum* genus. A) Densitree plot of 1,000 randomly sampled trees with the best posterior probabilities based on the concatenation of single-copy autosomal genes (green traces). Tips are labelled with scientific names of *Microbotryum* fungi and of their hosts. Boxed letters serve as a guide for species labels in panels D and E. Asterisks indicate the species whose genomes are reported here for the first time. Photos of diseased flowers by M.E. Hood. A) Rearrangement scenarios as cartoons and presence of evolutionary strata across the phylogenetic tree (vertical colored bars). Extant recombining regions in mating-type chromosomes are colored in blue or purple when corresponding to the ancestral HD or PR chromosome, respectively. Non-recombining regions are colored according to the linear gradient displayed at the top in millions of years since the onset of recombination suppression. Pale green chromosomes are autosomes derived from the fission of the long arm of the ancestral HD chromosome or the short arm of the ancestrral PR chromosome. C) Diagram of ancestral mating-type chromosomes showing the different regions involved in recombination suppression events. Regions are colored according to the number of independent suppression of recombination events in which these have been involved (gradient at the right). Abbreviations: distCEN, distal region to the HD centromere on the long arm; proxiCEN, proximal region to the HD centromere on the long arm; CEN:HD, region between the centromere and the HD locus; HD:short, region between the HD locus and the short arm telomere; short:CEN, short arm of the PR chromosome; CEN:PR, region between the centromere and the PR locus; proxiPR, region proximal to the PR locus towards the long arm telomere. D) Density distributions of date estimates for recombination suppression events. Note that the range between 0 and 0.2 million years ago (MYA) is expanded. The 22 time points used for studying the tempo of degeneration are numbered on the plots. E) Densitree plots for the different regions involved in recombination suppression events. For each region the number of genes indicated in the insets were concatenated to reconstruct 10^7^ ultrametric trees, from which 1,000 randomly sampled trees among the 7.5 x 10^6^ with best posterior probabilities are plotted. Trees are colored according to the named strata in *M. v. paradoxa* (Figure 2X). Shadowed clades show trans-specific polymorphism (i.e. alleles clustered by mating-type rather than by species). The species *M. v. paradoxa* (g) branches differently in the short arm of the PR chromosome and the PR:CEN regions than in the species tree. Note that the tree based on the short arm of the PR chromosome has two alternative topologies. Large dots on internal nodes correspond to the suppression of recombination events (colored red for those first described here). Symbols at the tips correspond to the alternate mating-types and they are colored according to the focal stratum.

In *M. v. melanantha* and *M. coronariae*, the mating-type chromosomes carried both PR and HD loci and were found to result from the fusion of the whole ancestral PR chromosome with the short arm of the ancestral HD chromosome (Figures 1A and S1), as was previously found in *M. lychnidis-dioicae*, *M. silenes-dioicae* and *M. violaceum sensu stricto* (Branco, et al. 2017; Branco, et al. 2018). Thus, the mating-type chromosomes of the five species in this clade resulted from the same chromosomal rearrangements. However, trans-specific polymorphism analyses, detailed below, suggested three independent events of complete recombination cessation in this clade (Figure 1E). Indeed, as soon as recombination cessation links a gene to the mating-type locus, the alleles of this gene associated to the alternative mating-type alleles start diverging and accumulate mutations independently. The alleles of such a non-mating-type gene will thus cluster per mating type, even across speciation events. The node to which trans-specific polymorphism extends therefore indicates the date of complete recombination cessation (Hartmann, et al. 2021). Despite the different rearrangements having led to HD and PR linkage across the *Microbotryum* phylogeny, a subset of 32 genes was shared among nearly all types of black strata, i.e. were initially located between the mating-type loci for all types of rearrangements except in *M. silenes-acaulis*. Genealogies for these 32 genes, including also their sequences from *M. silenes-acaulis* and *M. intermedium*, despite being in recombining regions in these species, revealed patterns of trans-specific polymorphism consistent with three independent events of complete recombination cessation (Figure 1E): one in *M. v. melanantha* (with no trans-specific polymorphism), one in the lineage leading to *M. lychnidis-dioicae* and *M. silenes-dioicae* (two species sharing trans-specific polymorphism), and one in the lineage leading to *M. coronariae* on *L. flos-cuculi* and *M. violaceum* (s.s.) on *S. nutans* (also sharing trans-specific polymorphism one with each other). Alternatively, there could have been a single chromosomal rearrangement event in the common ancestor of the clade encompassing these five species, but without initially complete recombination cessation, this occurring later, independently in the three lineages. For our purposes, studying the consequences of recombination suppression, the three independent events of complete recombination cessation are the important events, regardless of whether the initial chromosome fusion occurred once or three times independently, and we therefore considered in the following these three black evolutionary strata as independent.

In *M. v. tatarinowii*, the mating-type chromosomes resulted from the fusion of a part of the long arm of the PR chromosome with the short arm of the HD chromosome (Figure 1A; Figure S1). We found the remaining part of the ancestral PR chromosome on an autosomal contig in the two haploid genomes of *M. v. tatarinowii*. This represents the same type of chromosomal rearrangement leading to PR-HD linkage as previously found in *M. v. caroliniana* (Branco, et al. 2018). However, the phylogenetic placement of the two species shows that they represent independent events (Figure 1A). In addition, we improved the assemblies of *M. scabiosae* (Table S2) and refined the chromosomal rearrangement scenario for *M. v. caroliniana*. We could thus refine the breakpoint locations for the fissions and fusion generating their mating-type chromosomes (Figure S3).

### Evolutionary strata

We investigated the existence of evolutionary strata in *M. v. melanantha*, *M. v. tatarinowii* and *M. coronariae* following the methods used previously (Branco, et al. 2017; Branco, et al. 2018), *i.e*., by plotting the synonymous divergence calculated between the alleles from the genomes of opposite mating types along the ancestral-like gene order of *M. intermedium* (Figure 2). The rationale of this method is that, as soon as some genes become linked to the mating-type genes, the alleles associated with the two mating-type alleles gradually accumulate substitutions independently, so that their differentiation increases with the time since recombination suppression. Stepwise extension of recombination suppression therefore yields a pattern of discrete strata with different levels of differentiation between alleles on the mating-type chromosomes. In contrast, we expect high levels of homozygosity in these selfing species in recombining regions, and therefore no differentiation between the two genomes derived from a diploid individual, including in pseudo-autosomal regions. The evolutionary strata may however be difficult to distinguish using only differentiation levels, so we delimited strata based on: i) the distribution of recombination suppression across the phylogenetic tree to assess when recombination suppression evolved and thereby identifying distinct evolutionary strata, *i.e*. genomic regions that stopped recombining at different times (Figure 1D), as well as ii) the level of trans-specific polymorphism (Figure 1E), that is also a strong indicator of the time of recombination suppression.

**Figure 2:**
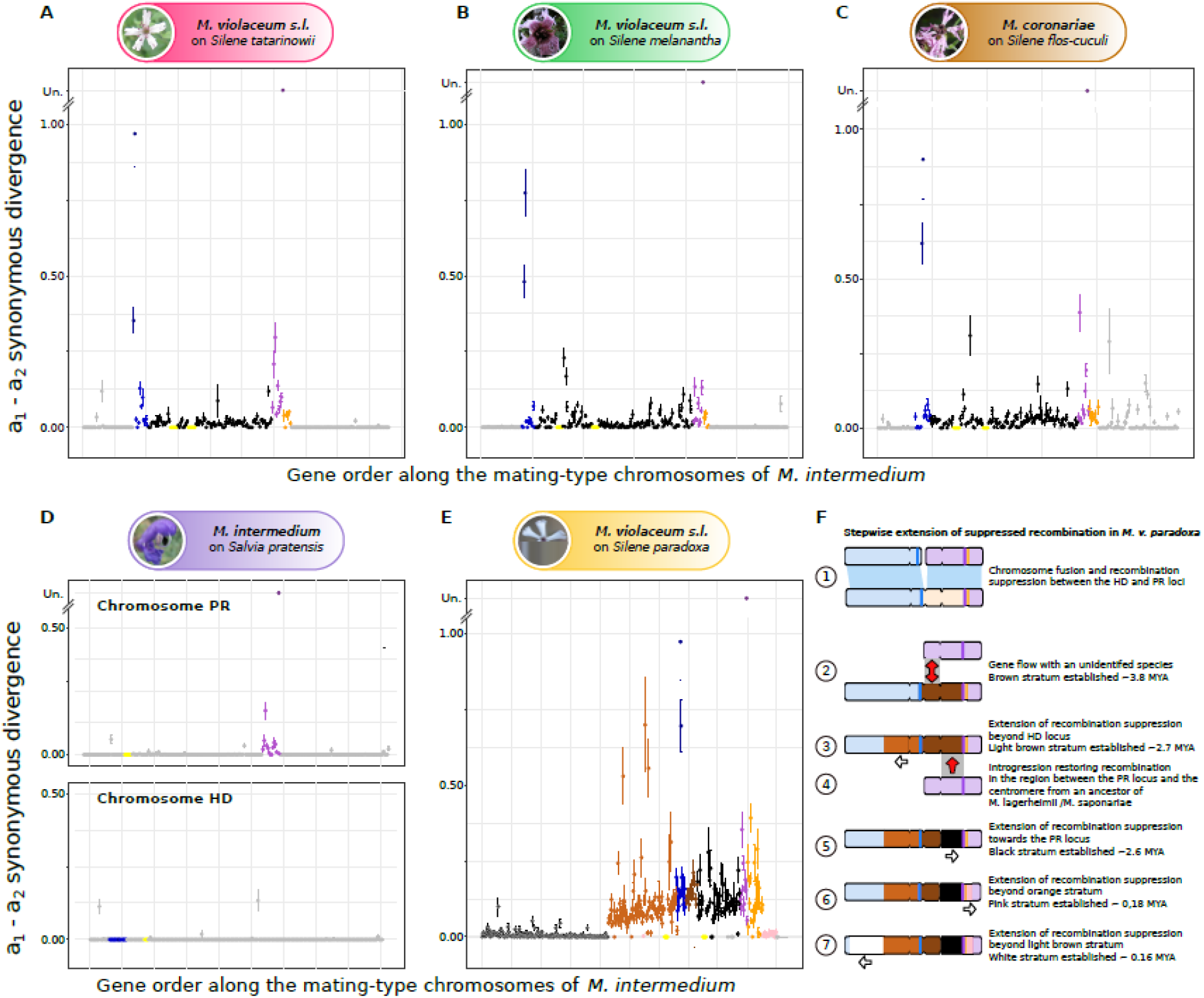
Allelic divergence analysis between mating-type chromosomes for the newly sequenced species and evolution of the mating-type chromosomes in *Microbotryum v. paradoxa*. For each species (A, *Microbotryum v. tatarinowii*; B, *M. v. melanantha*; C, *M. coronariae;* D, *M. intermedium* and; E, *Microbotryum v. paradoxa*), we show the synonymous divergence calculated between alternative mating types and plotted along the *M. intermedium* mating-type chromosome gene order. Zero d_S_ values between mating-type chromosomes indicate recombination in these selfing species and the level of d_S_ between mating-type chromosomes increases with time since recombination suppression. Different evolutionary strata, i.e., different steps extending recombination suppression, are indicated by different colours. Divergence between the a1 and a_2_ pheromone receptor (PR) alleles was too extensive for d_S_ to be calculated (depicted as “Un” for unalignable). The different strata were delimited based on their presence in the various species across the phylogenetic tree (Figures 1A and 1B) and their level of trans-specific polymorphism (Figure 1E). B. Scenario of the mating-type chromosome evolution in *Microbotryum v. paradoxa*.

We identified four evolutionary strata in each of *M. v. melanantha*, *M. v. tatarinowii* and *M. coronariae*. Three strata are ancient and shared among most *Microbotryum* species, *i.e*., the purple stratum around the PR locus, the blue stratum around the HD locus, and the orange stratum adjacent to the purple stratum (Figures 1A and 2A-D). In addition, recombination suppression that linked the HD and PR loci together evolved at different times in these three species (Figure 1D). We called all these regions involved in the different events of HD-PR linkage the black evolutionary strata, but they resulted from different types of chromosomal fissions and fusions and therefore encompass distinct gene sets (Figure 1A). We identified no additional evolutionary strata younger than the PR-HD linkage events in any of the species that we sequenced in the present study. For all statistical analyses we considered only independent events of recombination suppression, analysing the genomic regions corresponding to the species-specific strata or to the independent black strata (independent events linking HD and PR loci to each other or to their centromere that often trapped different gene sets). We did not include in our statistical analyses the oldest strata shared by multiple species that arose in a common ancestor, although we checked that similar results were obtained when including all strata.

The mating-type chromosome of *M. v. paradoxa* was formed by the fusion of the whole HD and PR ancestral mating-type chromosomes (Branco, et al. 2018). The two most recent strata in *M. v. paradoxa* identified previously are distal to the mating-type loci and were called pink and white strata (Figure 2E). The analysis of single-copy genes in its mating-type chromosomes showed that the recombination suppression previously called black stratum in *M. v. paradoxa* actually occurred in multiple steps. The oldest event of recombination suppression corresponded to the genes located ancestrally between the HD-chromosome centromere and the PR locus (“brown” stratum, Figure 2F), thus linking the two mating-type loci. Recombination later stopped distal to the ancestral HD chromosome centromere on the long arm of the ancient HD chromosome (“light-brown” stratum), spreading to non-mating-type genes (Figure 2F). This was followed by an introgression from an ancestor of *M. lagerheimii* and *M. saponariae*, with still-recombining mating-type chromosomes, thus restoring recombination between the centromere and the PR locus. Recombination was subsequently again suppressed in this introgressed region (“black” stratum of *M. v. paradoxa*, Figure 2F). The introgression and the independence and timing of the different events of recombination cessation are supported by i) the non-overlapping distributions of the estimated times of recombination suppression for the brown and black strata (Figure 1D), ii) the different levels of differentiation between mating-type chromosomes (Figure 2E) and iii) the different tree topologies for the genes in the brown or light-brown strata versus the black stratum (Figure 1E), as explained below. The light brown stratum may even be constituted itself of different strata or represent a gradual extension of recombination suppression, as its d_S_ values seem to decrease progressively when going away from the mating-type locus (Figure 2E).

The black in *M. v. paradoxa* indeed displayed a different placement of *M. v. paradoxa* compared to the species tree: *M. v. paradoxa* was sister species to the clade encompassing *M. coronarie*, *M. lychnidis-dioicae* and *M. violaceum s.s*. in the species tree (Figure 1A), while it was placed as a sister species to the *M. lagerheimii*-*M. saponariae* clade in the black stratum tree (Figure 1E). This discrepancy strongly supports an introgression event of the black stratum in *M. v. paradoxa* shortly after speciation, from the ancestor of *M. lagerheimii* and *M. saponariae* (Figure 2F). This reinforces the view that mating-type chromosomes in fungi may be particularly prone to introgression, as this process may counteract the effect of degeneration following recombination suppression (Corcoran, et al. 2016; Hartmann, et al. 2020; Sun, et al. 2012). The black stratum in the ancestor of *M. v. paradoxa* has indeed been introgressed from the ancestor of *M. lagerheimii* and *M. saponariae* that had likely not yet stopped recombining in these regions, as their recombination suppression events in this region is younger than their speciation (Carpentier, et al. 2019). This introgression from recombining regions has likely temporarily restored recombination in these regions in the *M. v. paradoxa* lineage, explaining the younger recombination suppression date and the presence of the introgression on both *M. v. paradoxa* mating-type chromosomes. The introgression despite recombination suppression in the *M. v. paradoxa* lineage could have occurred via gene conversion and/or because the recombination was not suppressed with the recombining chromosomes of the ancestor of *M. lagerheimii* and *M. saponariae*. The brown stratum of *M. v. paradoxa* also had a different placement than the species tree and from the black stratum (Figure 1E), possibly resulting from an introgression from an unidentified species.

### Absolute dating of recombination suppression events

We estimated the dates when recombination was suppressed for the 22 strata for which we had at least ten genes or 10,000 aligned codons among single-copy genes in non-recombining regions. As soon as a gene is permanently linked to the mating-type loci, its alleles will independently accumulate substitutions that will remain associated with their linked mating-type allele. We therefore inferred the age of recombination suppression by estimating the divergence time between the alleles associated with the a_1_ and a_2_ mating types, using Beast v2.5.0. We used the split of *M. lychnidis-dioicae* and *M. silenes-dioicae* at 0.42 MYA as a calibration point (Gladieux, et al. 2011). The estimated times ranged from 3.6 to 0.15 million years before present for the various independent black strata (Table S3). We estimated divergence times ranging from 3.82 to 0.15 million years in *M. v. paradoxa* strata and in the range of 0.07 to 0.25 million years for the young strata such as the light-blue stratum of *M. v. caroliniana* and the red stratum of *M. lychnidis-dioicae* and *M. silenes-dioicae* (Table S3).

### Identification of optimal codons

To identify optimal codons in *Microbotryum* fungi, we investigated which codons were preferentially used in highly expressed autosomal genes in the two distantly related species, *M. lychnidis-dioicae* and *M. intermedium* (Figure 1). Codon usage bias is expected to be strongest for the most highly expressed genes and selection for optimal codon use to be more effective on autosomes because recombination occurs. We had expression data for haploid cells cultivated clonally on nutritive medium (Table S4), for the two mating types separately. To study autosomal codon usage, we performed a within-group correspondence analysis (WCA; (Charif, et al. 2005)), *i.e*., a multivariate statistical method summarizing codon count data, normalizing the count of each codon by the average count of all the codons encoding for the same amino acid (Suzuki, et al. 2008). For the *M. intermedium* a_1_ mating type, the first principal component axis summarized 16.55% of the codon usage variation (Figure S4) and was significantly negatively correlated with the gene expression level (Table S5). Therefore, the genes with the lowest coordinates on the first principal component axis were those most highly expressed, where we expect strongest bias towards optimal codons. We performed chi-square tests to compare the codon counts per amino acid of the genes with the 10% highest coordinates and the 10% lowest coordinates on the first axis. For each amino acid, we inferred the optimal codon to be the one with the highest adjusted standardized residuals (*i.e*., largest counts compared to expected under the hypothesis of random use of synonymous codons) in highly expressed genes from chi-square tests (Table 1). We inferred the same optimal codons when using the autosomes from the a_2_ mating type of *M. intermedium* or from either of the two *M. lychnidis-dioicae* mating types (Table S6). Because *M. intermedium* and *M. lychnidis-dioicae* are very distantly related species in our dataset (last common ancestor at ca. 20 million years before present, Figure 1A), and share the same optimal codons, we assumed the optimal codons to be the same in all other *Microbotryum* species. The optimal codons were more GC rich than the other synonymous codons (Table 1).

**Table 1:**
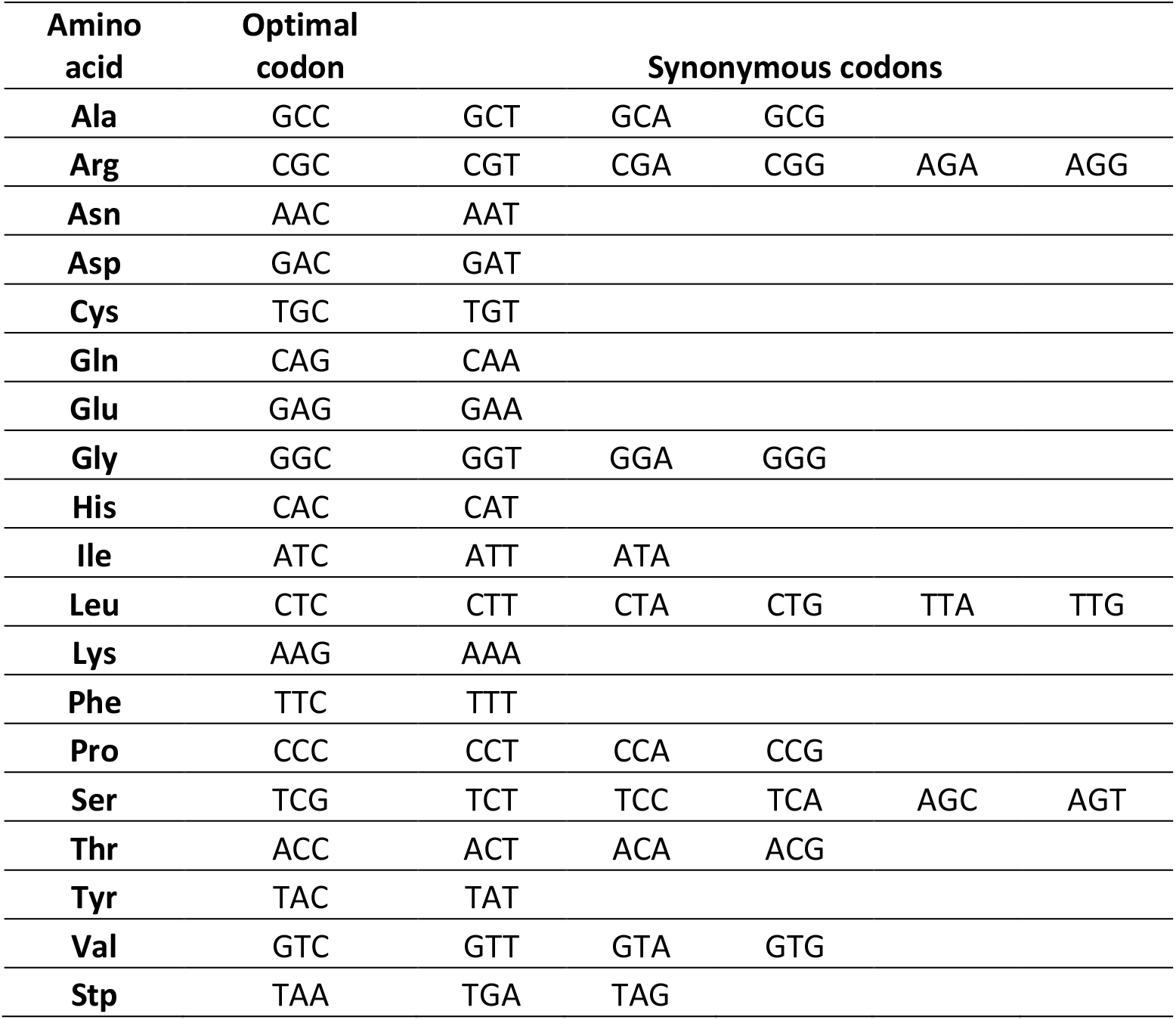
Optimal codons in *Microbotryum*.

| Amino acid | Optimal codon | Synonymous codons |  |  |  |  |
| --- | --- | --- | --- | --- | --- | --- |
| **Ala** | GCC | GCT | GCA | GCG |  |  |
| **Arg** | CGC | CGT | CGA | CGG | AGA | AGG |
| **Asn** | AAC | AAT |  |  |  |  |
| **Asp** | GAC | GAT |  |  |  |  |
| **Cys** | TGC | TGT |  |  |  |  |
| **Gln** | CAG | CAA |  |  |  |  |
| **Glu** | GAG | GAA |  |  |  |  |
| **Gly** | GGC | GGT | GGA | GGG |  |  |
| **His** | CAC | CAT |  |  |  |  |
| **Ile** | ATC | ATT | ATA |  |  |  |
| **Leu** | CTC | CTT | CTA | CTG | TTA | TTG |
| **Lys** | AAG | AAA |  |  |  |  |
| **Phe** | TTC | TTT |  |  |  |  |
| **Pro** | CCC | CCT | CCA | CCG |  |  |
| **Ser** | TCG | TCT | TCC | TCA | AGC | AGT |
| **Thr** | ACC | ACT | ACA | ACG |  |  |
| **Tyr** | TAC | TAT |  |  |  |  |
| **Val** | GTC | GTT | GTA | GTG |  |  |
| **Stp** | TAA | TGA | TAG |  |  |  |

### Decrease in optimal codon frequency in non-recombining regions

We determined whether each codon was optimal or not. We did this for each haploid genome of the 13 *Microbotryum* species included in this study, separately for on the autosomes and the different compartments of the mating-type chromosomes, *i.e*. the recombining pseudo-autosomal regions (PARs) and distinct evolutionary strata with different ages of recombination suppression. However, several of these evolutionary strata are small and contain few genes, which precludes a reliable estimate of optimal codon usage. We therefore only considered the 22 largest non-recombining regions (with at least 19 genes; Figure 1). This was to test whether the frequencies of optimal codons differed between genes in these non-recombining regions and in autosomes. However, because we found that optimal codons were enriched in GC in *Microbotryum*, a decrease in the frequency of optimal codons in the non-recombining regions compared to recombining regions could not only be due to a lower efficacy of selection for optimal codon usage but also a reduction in GC-biased gene conversion in non-recombining regions. It should be possible to distinguish these two effects because GC-biased gene conversion affects both coding and non-coding sequences while lower efficacy of selection on codons should only impact coding sequences (i.e. on exons but not on introns). We therefore introduced the difference in GC content between coding sequences (GC^exon^) and introns (GC^intron^) per gene as a covariate in the analyses of optimal codons as an estimation of the relative effects of GC-biased gene conversion versus selection on codon usage. We performed a logistic ANCOVA for each genome to explain the variation in the status of codons (*i.e*., optimal versus non-optimal) by the genomic compartment, by the difference in GC content between coding sequences (GC^exon^) and introns (GC^intron^), and by their interactions (see Figure S5 for raw data distribution). These factors allowed us to test (i) whether the optimal codon usage varied significantly among the different genomic compartments, (ii) whether the optimal codons usage varied with the difference in GC content between coding sequences and introns (GC^exon^ - GC^intron^), which estimates the impact of GC-biased gene conversion, and (iii) whether the strength or direction of the relationship between optimal codon use and GC-biased gene conversion varied among the different genomic compartments. We wanted to assess whether codons were less often optimal in non-recombining regions (tested through the genomic compartment effect), and to assess whether this could be adequately explained solely by a lower frequency of GC-biased conversion in non-recombining regions, which was tested through the interaction between the genomic compartment and the quantitative measure of the effect of GC-biased conversion.

We found few significant differences in optimal codon usage between autosomes and the recombining regions located on mating-type chromosomes (Table S7a). In contrast, across all species and mating types, codons were significantly less often optimal in non-recombining regions than in autosomes (Table S7a, Figure 3A, Figures S6 and S7), except for the very recently evolved non-recombining region of the a_2_ HD *M. lagerheimii* mating-type chromosome (Table S7a). Codons were thus less often optimal in older non-recombining evolutionary strata than in recombining regions. To understand whether this was a signal of degeneration due to the lower efficacy of purifying selection in non-recombining regions, we needed to ascertain whether the lower frequency of optimal codons could be explained simply by a lower frequency of recombination-mediated gene conversion, gene conversion being typically GC-biased and optimal codons being, on average, more GC rich than less optimal codons. To test for this, we used the difference in GC^exon^ - GC^intron^ as a covariate. Because low efficacy of selection on codon usage should influence only coding sequences while GC-biased conversion should affect all sequences, large excesses in GC^exon^ compared to GC^intron^ points to the predominant action of selection whereas no difference points to the predominant action of GC-biased gene conversion. Across the genomic landscape of autosomes, genes that showed low difference in GC content between exons and introns were most rich in optimal codons, indicative of stronger influence of GC-biased conversion. As the difference in GC content between exons and introns increased the frequency of optimal codons declined. This was the case for all the genomes (Table S7a; Figure 3B), except the a_1_ genomes of *M. lychnidis-dioicae*, *M. v. melanantha* and *M. violaceum* (*s.s*.) for which the effect was not significant (Table S7a). This suggests that GC-biased gene conversion, in addition to selection for optimal codon usage, increases the frequency of optimal codons in autosomes. Indeed, selection for optimal codons is strong only in highly expressed genes, while GC-biased conversion can act on all genes. This also means that the generally lower frequency of optimal codons in non-recombining regions could in principle be due to less frequent GC-biased gene conversion in these regions compared to autosomes.

**Figure 3:**
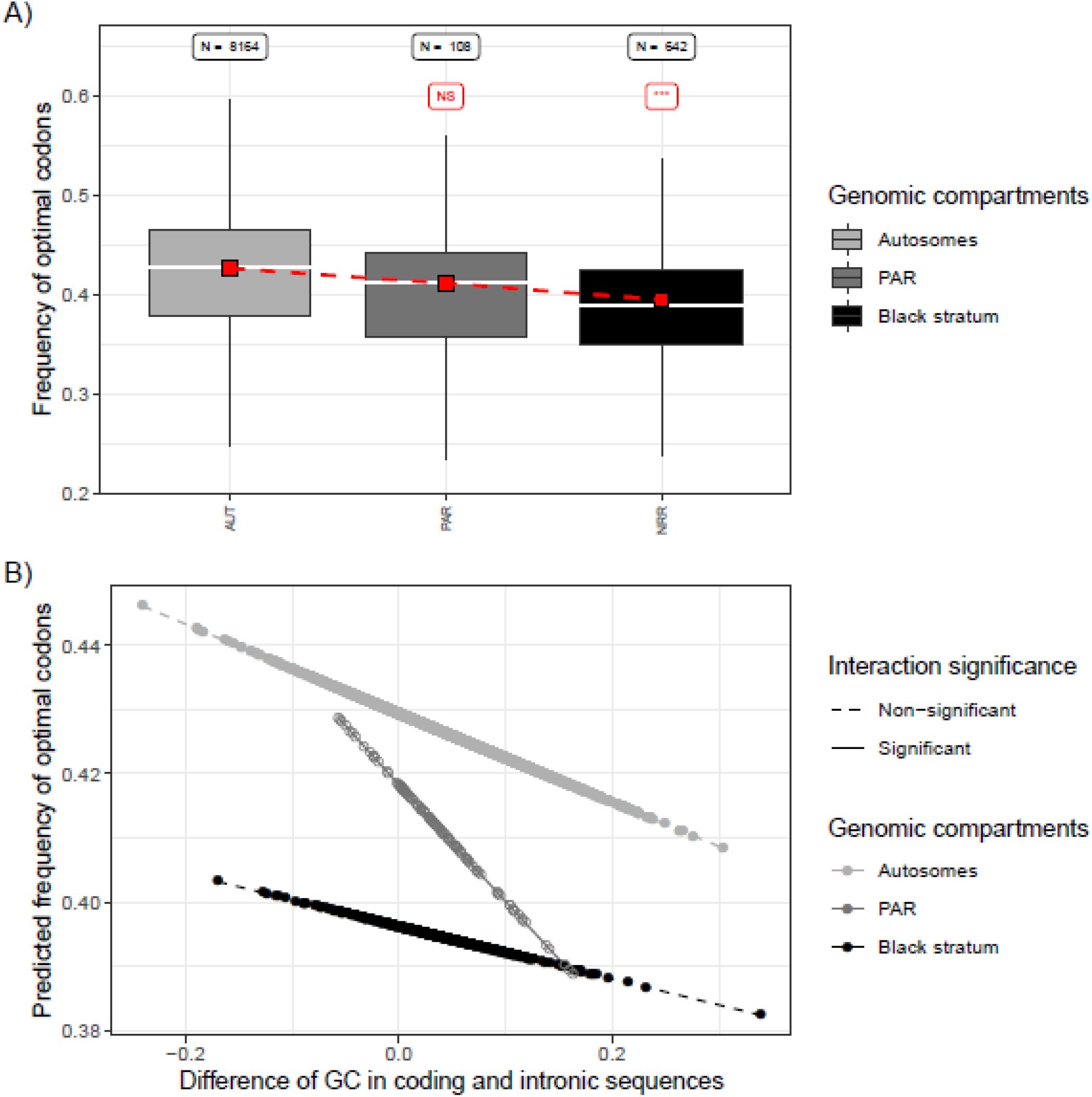
Analysis of the frequency of optimal codons in the a_1_ genome of *Microbotryum silenes-dioicae*. A) Distribution of the frequency of optimal codons across genomic compartments. Boxplot grey shade refers to the genomic compartment. For each genomic compartment, the red square indicates the mean value of the predicted frequency of optimal codons, obtained from the logistic regression explaining the frequency of optimal codons by the genomic compartment, by the difference in GC content between coding (GC^exon^) and intronic (GC^intron^) sequences and their interaction. The sample size (N) is labelled on top of the corresponding boxplot, as well as the significance level of the difference between a given genomic compartment and the autosomes in red (NS: non-significant; ***: p < 0.001) from the logistic regression. B) Frequency of optimal codons predicted by the logistic regression plotted against the difference in GC content between coding and intronic sequences. From the statistical analysis, significant differences in slope between a genomic compartment and the autosomes are indicated by solid line and filled dots, while non-significant interactions are indicated by dotted line and open dots. Similar analyses for all species studied here are presented in Figures S6 and S7 in the supplementary materials.

However, the relationship between the difference in GC^exon^ and GC^intron^ and optimum codon usage rarely differed between autosomes and non-recombining regions, most interactions being non-significant (Table S7a; Figure 3B). In most genomes, we indeed observed the same slope of optimal codon frequency as a function of the difference in GC^exon^ and GC^intron^ in non-recombining regions as in autosomes (Figures 3B and S6). This means that the codons were not only less optimal in non-recombining regions in the genomic areas where GC-biased conversion had the highest impact (low difference in GC^exon^ and GC^intron^, left Figure 3B) but also where it had little influence, i.e., where only selection acts (high difference in GC^exon^ and GC^intron^, right Figure 3B). Such similar slopes for autosomes and non-recombining regions, together with lower overall values of optimal codon frequency in these non-recombining regions, indicate that selection for optimal codons has a lower efficacy across the genomic landscape in non-recombining regions and that the GC-biased conversion has similar impact in recombining and non-recombining regions. We also performed the same statistics using all evolutionary strata, even the shortest ones, and we obtained similar conclusions as when using only the largest ones (Figure S8, Table S7b).

### Control for ancestral gene expression level

Selection on codon usage of genes could be weaker in non-recombining regions if gene expression in these regions had been, by chance, low in these genomic regions even before recombination was suppressed. To investigate whether genomic compartments differed in the average gene expression level before recombination suppression, we inferred the ancestral expression level of each gene from the expression level of its ortholog in *M. intermedium*, which has no recombination suppression on its mating-type chromosomes except in small regions around the mating-type loci themselves. Of course, some gene-specific expression levels may have changed in *M. intermedium* since its divergence from the last common ancestor, but, without recombination suppression in this species, there has likely been no major change in gene expression across large genomic regions since the last common ancestor of the studied *Microbotryum* species. That gene expression level changed little across the *Microbotryum* genus in recombining regions was supported by highly significant correlations of gene expression levels of shared autosomal genes measured in similar culture conditions between *M. intermedium* and *M. lychnidis-dioicae* (Tables S8 and S9).

For each of the two haploid genomes across the 13 species, we compared the mean and variance of the inferred ancestral gene expression level (*i.e*., the gene expression level in *M. intermedium*) between the current genomic compartments using a Kruskal-Wallis’ test and a Levene’s test, respectively. For each species we tested for differences in the inferred ancestral gene expression between the current genomic compartments, *i.e*., autosomes, PARs and the various evolutionary strata that lack recombination. We found no significant difference in mean or variance of the inferred ancestral gene expression level between the current recombining and non-recombining regions for any genome (Table S10). Therefore, differences in frequencies of optimal codons across the genomic compartments are unlikely to be explained by differences in ancestral gene expression level between genomic compartments.

### Tempo of degeneration in codon usage and in protein-coding genes

The multiple independent events of recombination suppression in the *Microbotryum* genus provides an excellent case study for investigating the tempo of degeneration, allowing analysis of the relationship between degeneration level and time in non-recombining regions using independent data points. Here, we investigated the tempo of degeneration using the genes present in the regions with independent recombination suppression events between PR and HD loci, as well as the large species-specific strata. We excluded the shared strata common to most species because they do not provide independent data points (purple, blue and orange strata) and those that contain insufficient numbers of genes to ensure reliable estimates of the time since recombination suppression (e.g., the green strata in *M. lychnidis-dioicae*). For the black strata shared by multiple species (Figure 1), we took the mean value of variables for species descendent from the same recombination suppression event to avoid pseudo-replication. In total, we thus considered 22 evolutionary strata for these analyses (Table S11).

We tested whether the odds for a codon to be optimal (N^optimal^ / N^non-optimal^), the non-synonymous differences between alleles on the two mating-type chromosomes (d_N_) and the ratio of non-synonymous versus synonymous substitution rates (d_N_/d_S_) for the genes located in the non-recombining regions covaried with the time since recombination suppression, including species as a cofactor and, for optimal-codon analyses, ancestral gene expression level as a covariable. We tested linear, logarithmic or quadratic relationships with the time since recombination suppression on the one hand and d_N_, d_N_/d_S_ or the odd for a codon to be optimal on the other hand.

The odds for a codon to be optimal was best explained by considering the joint effect of its ancestral expression level, the species and a quadratic effect of the age since recombination suppression (Figure 4a, Table S12). This means that the decrease in the frequency of optimal codons slowed with time, reaching an asymptote at ca. 3MY (Figure 4a). As expected, the odds for a codon to be optimal increased with its ancestral expression level (Figure S9; Table S12). The asymptote for the frequency of optimal codons seemed at a higher value than expected under random codon usage. There are indeed 18 amino-acids encoded by more than one codon and 59 codons in total discarding the three stop codons and the two single-codon aminoacids; this gives a frequency of optimal codons of 18/59=0.30 under random usage while the asymptote is at ca. 0.43 and the lowest frequencies at 0.40 (Figure 4a).

**Figure 4:**
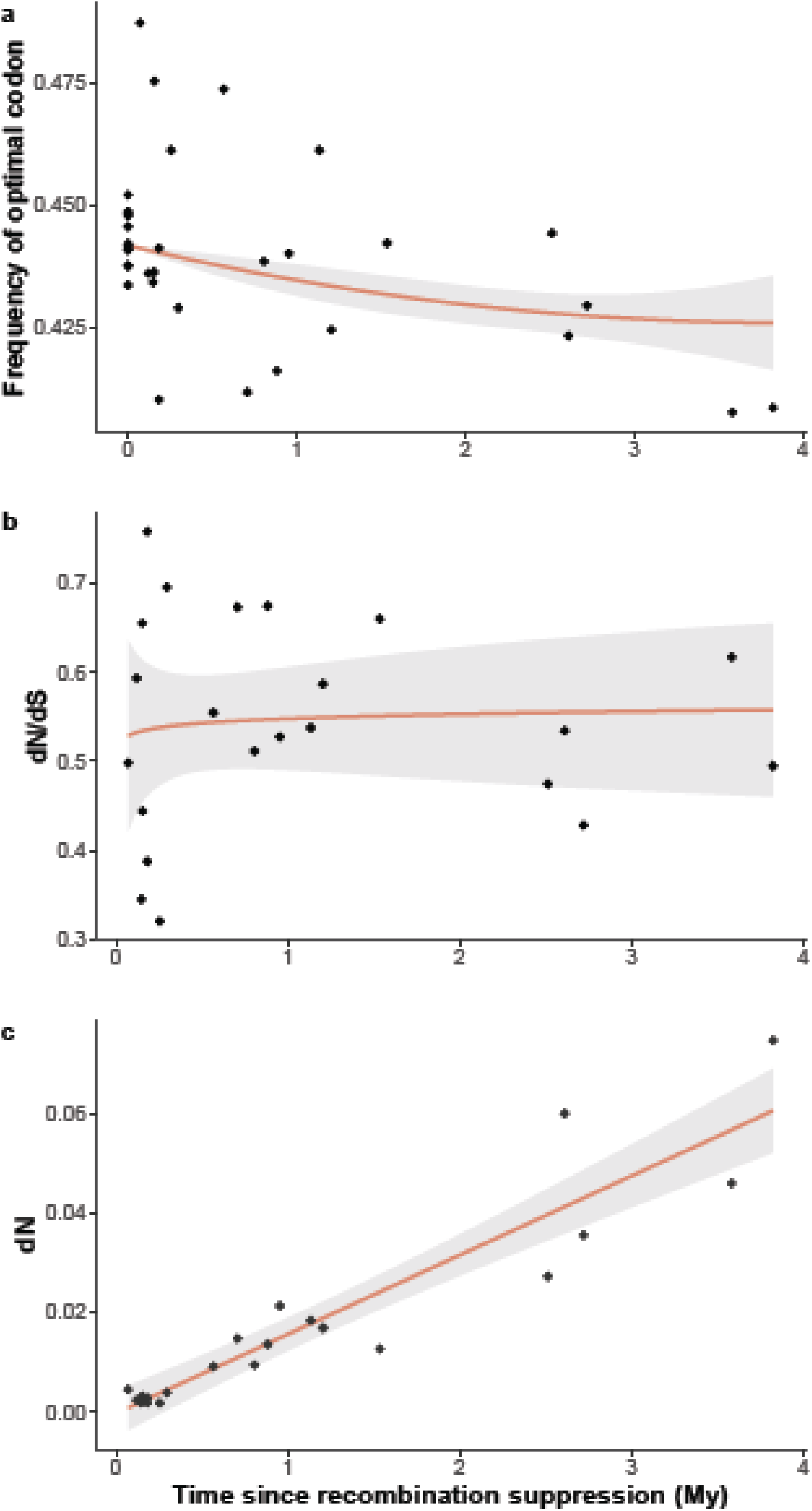
Tempo of degeneration in *Microbotryum* fungi. A) Frequency of optimal codons N^optimal^ / (N^optimal^ + N^non-optimal^) as a function of time since recombination suppression (in million years). Note that the statistical analyses were performed on the the odds of optimal codons (N^optimal^ / N^non-optimal^) and not the frequency as plotted here. B) Non-synonymous over synonymous divergence (d_N_/d_S_) between alleles on the two mating-type chromosomes, as a proxy of the strength of purifying selection, as a function of time since recombination suppression (in million years). C) Accumulation of non-synonymous changes accumulation (d_N_) between alleles on the two mating-type chromosomes as a function of time since recombination suppression (in million years). In all panels, the dots represent the data. Predictions from the models are show as red lines and the 95% confidence intervals of the prediction as grey areas.

The d_N_/d_S_ ratio rapidly increased following recombination suppression and was best explained by the log of time since recombination suppression (Figure 4b, Table S12). However, the d_N_/d_S_ ratio very rapidly reached a plateau, actually remaining almost constant at a value around 0.55. This means that purifying selection decreased very rapidly following recombination suppression to a constant level, intermediate between strong purifying selection (d_N_/d_S_=0) and neutral evolution (d_N_/d_S_=1). Accordingly, non-synonymous differences between mating-type chromosomes (d_N_) increased linearly with stratum age, at a rate of 0.015 per MY (Figure 4c). In contrast, both d_N_ and d_S_ were near zero in recombining regions between alleles from haploid cultures of opposite mating types deriving from a diploid individual (Figure 5), these fungi being highly homozygous due to high selfing rates.

**Figure 5:**
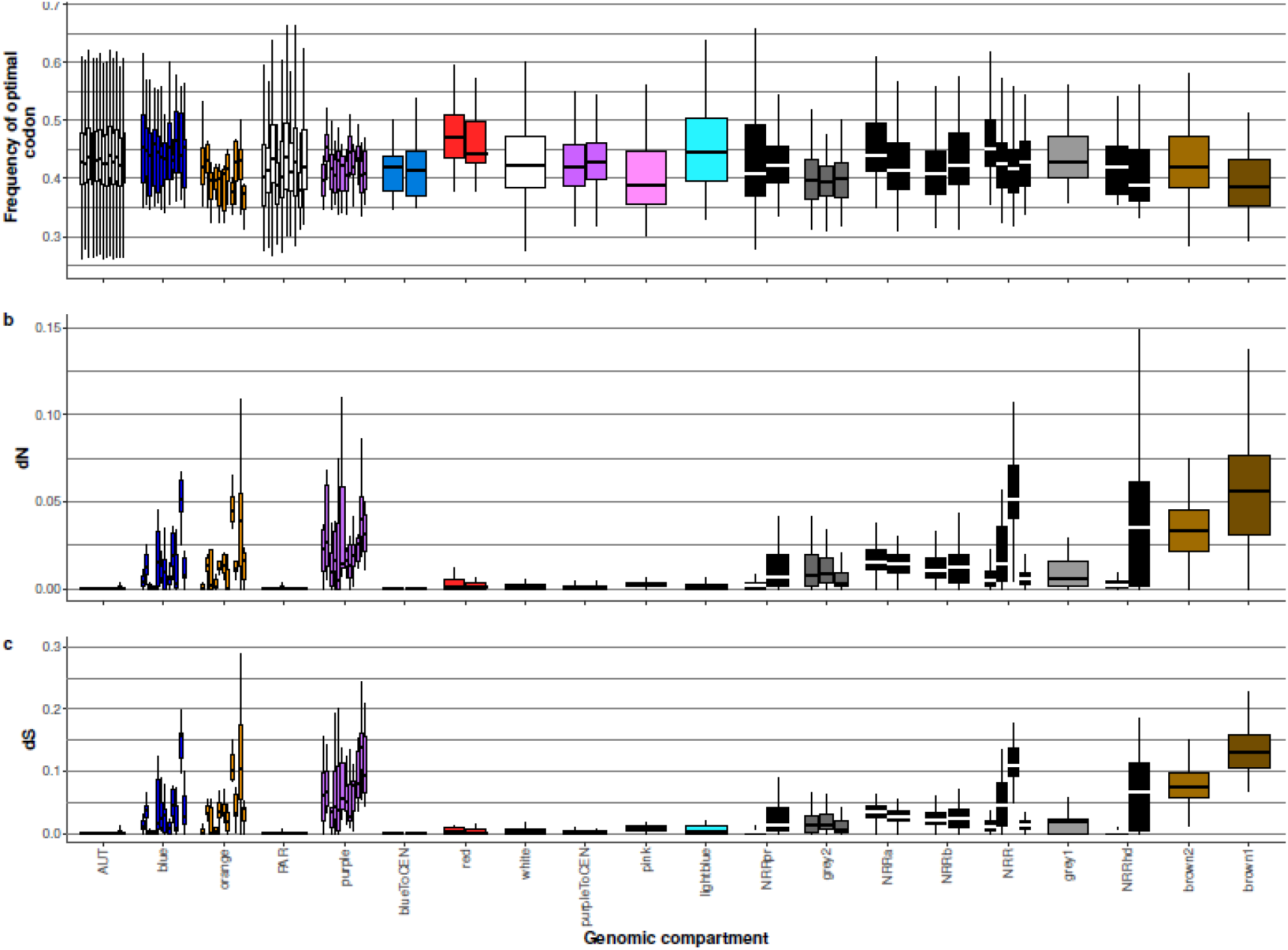
Degeneration in *Microbotryum* fungi, with values for all genomic compartments and species. Boxplots of frequencies of optimal codons (A) and non-synonymous differences (d_N_) between alleles of alternative mating-type chromosomes (B) in all species and all genomic compartments. The X axis legend correspond to the one in Figure 1C.

For the degeneration tempo analysis, we used one single point per recombination suppression event to avoid pseudoreplication and we excluded the small strata with too few genes to provide reliable age estimates. Plotting the values for all strata and all species (Figures 5 and S8) showed similar degeneration levels among species in shared strata and thus repeatability of degeneration; this further supports the degeneration tempo analyses above. Figures 4 and 5 also show that the frequencies of optimal codons in recombining regions (autosomes and PAR) were high and relatively homogeneous among species.

## Discussion

This study takes advantage of multiple events of recombination suppression on the mating-type chromosomes of closely related fungal species to document the tempo of genomic degeneration in non-recombining regions (*i.e*., the pace at which deleterious mutations accumulate over time). We quantified the rate of degeneration by estimating the age of each non-recombining region, *i.e*., the time since recombination suppression, and by examining how the degree of degeneration estimated using two different measures, i.e., non-synonymous substitution accumulation and the reduction in optimal codon usage, varied with age. Few studies have focused on the degeneration of codon usage. This study further proposes a method for disentangling the effect of less efficient selection from other factors that influence codon usage in non-recombining regions. Indeed, codon usage is likely to be affected by ancestral gene expression level and GC-biased gene conversion, so studies of the consequences of relaxed selection on codon usage in non-recombining regions need to control for these two processes. We found that genes in non-recombining regions had fewer optimal codons than those in recombining regions. Controlling for levels of ancestral gene expression and GC-biased gene conversion allowed us to conclude that this was due to a decrease in selection efficacy in non-recombining regions. Furthermore, by examining independent non-recombining regions of varying age, we observed a rapid initial decrease in the frequency of optimal codon usage that then slowed to reach an asymptote. Non-synonymous substitutions, in contrast, accumulated in a nearly linear manner over the last 4 million years of suppressed recombination on *Microbotryum* mating-type chromosomes, in agreement with a steady level of purifying selection, intermediate between neutral evolution and strong selection against non-synonymous changes (d_N_/d_S_ = 0.55).

Multiple independent events of recombination suppression involving mating-type loci have been reported in anther-smut fungi (Branco, et al. 2017; Branco, et al. 2018; Carpentier, et al. 2019; Duhamel, et al. 2022). By sequencing and analysing the genomes of three additional *Microbotryum* species, we document three new independent events of mating-type chromosome rearrangements and fusions leading to PR-HD linkage, amounting to a total of eight such independent PR-HD linkage events in our present dataset. Linking mating-type loci is beneficial under selfing (Branco, et al. 2017; Nieuwenhuis, et al. 2013). The high number of independent events of PR-HD linkage through distinct genomic rearrangements shows the power of natural selection to generate repeated convergent phenotypes by independent pathways. In addition, there have been multiple extension events of the region of recombination suppression, at different times and in a stepwise manner in *Microbotryum* species. Here, we document additional previously unidentified evolutionary strata. In total, we analysed 22 genomic regions without recombination, each corresponding to an independent event of recombination suppression, providing the statistical power for a formal comparative study of genomic degeneration. Similar comparative analyses have been previously impossible because of the lack of such multiple independent events of recombination suppression across closely related species. The *Microbotryum* genus contains more than a hundred species (Hood, et al. 2010; Lutz, et al. 2008), so there are probably many as yet unexploited resources of genomic diversity that will allow deeper exploration of the effects of suppressed recombination in the vicinity of reproductive compatibility loci on genome evolution and in particular on the tempo of degeneration. The *Microbotryum* genus thus constitutes a unique model for understanding major evolutionary processes affecting the genomic structure of sexual eukaryotes.

We found that optimal codons were less frequently used in non-recombining regions compared to autosomes across multiple *Microbotryum* species. We interpreted this as a feature of degeneration due to reduced efficacy of selection in non-recombining regions, by controlling for possible confounding factors, *i.e*., lower GC-biased conversion or lower ancestral gene expression in non-recombining regions. Indeed, because optimal codons in *Microbotryum* are GC-rich, less GC-biased gene conversion in non-recombining regions compared to autosomes could lower the frequency of optimal codons in such regions. To test for this alternative explanation, we generated a covariate that estimated the strength of GC-biased gene conversion across genes in the genomes. We found similarly lower frequencies of optimal codons in non-recombining regions regardless of whether GC-biased conversion was high or low across the genomic landscape, and thus even when only selection acted. Thus, we can conclude that lower efficacy of selection contributes to the decrease in the odds of optimal codons in non-recombining regions in *Microbotryum* mating-type chromosomes. To our knowledge, this study constitutes the first attempt to analyse codon usage by controlling for confounding processes in statistical analyses.

We also checked that the differences in codon usage between autosomes and non-recombining regions were not due to differences in gene expression levels between these regions before the evolution of recombination suppression, as gene expression level affects codon usage (Novoa and Ribas de Pouplana 2012; Zhou, et al. 2016). Indeed, if expression levels were ancestrally lower for genes now located in non-recombining regions, selection on codon usage would have been weaker than in other autosomal regions, and lower frequencies of optimal codons would have been expected, regardless of the effect of recombination suppression. However, we found no difference in the ancestral expression levels between autosomal genes and those in current non-recombining regions. Altogether, we thus found robust evidence showing that lower efficacy of selection due to recombination suppression resulted in less optimal codon usage in *Microbotryum* species, as has been suggested in the non-recombining mating-type chromosome of the fungus *Neurospora tetrasperma* (Whittle, et al. 2011).

It has often been assumed that only weak selection would act on codon usage (Li, et al. 2009; Machado, et al. 2020), because synonymous substitutions should not greatly impact phenotypes. Contradicting this view, several studies, focusing at the gene level, have shown substantial impacts of synonymous substitutions on phenotypes (Carlini 2004; Lampson, et al. 2013; Machado, et al. 2020), sometimes leading to strong selection on codon usage (Machado, et al. 2020). In D*rosophila* species, codon usage was shown to be quite stable over long time frames with 11/12 species having the same preferred codons except those coding for serine (Vicario, et al. 2007). In the M*icrobotryum* genus, optimal codons were the same in *M. intermedium* and *M. lychnidis-dioicae*, two species having their last common ancestor at the basis of the studied *Microbotryum* clade, ca. 20 million years ago. The same optimal codons have therefore been maintained for several million years, suggesting strong and consistent selection. The preferred codons in two distantly related plant species, in different families, *S. latifolia* and *Arabidopsis thaliana*, are almost identical, despite their long divergence time (Qiu, et al. 2011). S*ilene latifolia* shows no decrease in optimal codon frequencies in non-recombining regions of the Y-chromosome (Bartolomé and Charlesworth 2006). In D*rosophila* in contrast, the rate of accumulation of unpreferred synonymous mutations was higher for the neo-Y chromosome than for the neo-X chromosome even for highly expressed genes (Bartolomé and Charlesworth 2006). Codon usage in non-recombining regions therefore requires more studies before we can draw generalities about its evolution.

Degeneration of protein-coding sequences in non-recombining regions of sex chromosomes has been much more studied than codon usage and has been consistently found in a variety of organisms, from animals and fungi to plants (e.g., (Bachtrog 2013; Bergero and Charlesworth 2011; Chibalina and Filatov 2011; Fontanillas, et al. 2015; Soojin and Charlesworth 2000; Whittle, et al. 2011). Overall trends were inferred from observations that older evolutionary strata, *e.g*., on sex chromosomes of papaya and supergenes in butterflies (Jay, et al. 2021a; Wu and Moore 2015) had higher non-synonymous substitution rates than younger strata. However, though theoretical models made predictions about the tempo of gene losses, empirical studies on the tempo of sequence degeneration were lacking (Bachtrog 2008), because data from sufficient numbers of independent recombination suppression events to examine the tempo of non-synonymous mutation accumulation did not exist.

The existence of multiple independent events of recombination suppression in *Microbotryum* fungi allowed reliable estimations of the tempo of degeneration in protein-coding sequences. The decrease in the frequency of optimal codons was rapid following recombination suppression and reached an asymptote at ca. 3My. The asymptote seemed above the value of the frequency of optimal codons expected at random, which suggests that selection is still acting. The d_N_/d_S_ ratio, a proxy for the degree of relaxed selection, very rapidly (within less than 0.25 MY) reached a constant value, remaining around 0.55. The efficacy of purifying selection thus very rapidly decreased after the onset of recombination suppression and then remained steady at the same intermediate value between strong purifying selection (d_N_/d_S_=0) and neutral evolution (d_N_/d_S_=1). Accordingly, non-synonymous differences (d_N_) increased linearly with stratum age, at a rate of 0.015 per MY. One could have expected that the strength of purifying selection would increase with time (*i.e*., d_N_/d_S_ would decrease with time) if accumulating more non-synonymous changes would have increasing deleterious effects, *i.e*., if fitness decreased more rapidly than linearly with the number of non-synonymous substitutions. One could have expected in contrast that the strength of purifying selection would decrease with time (*i.e*., d_N_/d_S_ would increase with time), if some gene copies became completely non-functional. This may be the case, for example, if selection on their function occurs primarily in the diploid (dikaryotic) stage such that deleterious mutation can remain sheltered under the permanent heterozygosity afforded by recombination suppression. We did not observe such non-linear effects here, but they may occur at longer time scales following recombination suppression. In addition to revealing the tempo of degeneration, this is the first study to our knowledge that disentangles the effects of reduced selection efficacy and GC-biased gene conversion on the frequency of optimal codons and that investigates the tempo of degeneration using multiple independents events of recombination suppression.

We used synonymous and non-synonymous substitutions to estimate the time of divergence between sequences, as is typically done. We analysed how d_N_, d_N_/d_S_ and the odds of optimal codons evolved over time since recombination suppression. These different measures depend on substitution rates, with the influence of both mutation and selection, and it is precisely this effect of selection we aimed to analyse. The codon usage and d_N_ should be strongly correlated to the total synonymous divergence only in the absence of purifying selection, which is precisely what we are studying. If selection is very efficient, optimal codon frequencies should not change and d_N_ should remain very low, as is in fact observed on autosomes, in contrast to the non-recombining regions of the mating-type chromosomes. We studied here how rapidly deleterious substitutions accumulate over time since recombination suppression, i.e., the shape of the curve of accumulation of deleterious mutations with time.

Understanding the tempo of degeneration is important for our knowledge on genomic evolution, on the origin of genetic diseases, and on the maintenance of regions without recombination. For example, it is important to note that the seven youngest evolutionary strata used in this study of degeneration tempo are still collinear between mating types (Branco, et al. 2017; Branco, et al. 2018; Carpentier, et al. 2019) and correspond to low levels of degeneration. In contrast, the evolutionary strata with strong signs of degeneration are also highly rearranged between mating types (Branco, et al. 2017; Branco, et al. 2018; Carpentier, et al. 2019). This implies in particular that the hypothesis postulating that inversions linked to a mating-type locus or a sex-determining region could easily reverse when they accumulate too much load (Lenormand and Roze 2021)likely does not hold, as the region would then have already accumulated further rearrangements, preventing reversion to the original state that would allow recombination. This would in turn support the hypothesis that the extension of recombination suppression could then simply be the result of the selection of recombination cessation for sheltering deleterious alleles (Jay, et al. 2021b).

### Data access

The dataset(s) supporting the conclusions of this article is(are) available in the [repository name] repository, [unique persistent identifier and hyperlink to dataset(s) in http://format]-will be added upon manuscript acceptance.

## Material and methods

### Genome sequencing

We sequenced haploid genomes of opposite mating types for one diploid strain of each of the following species: *M. v. tatarinowii* parasitizing *S. tatarinowii* (#1400, GPS 39°59′45.4″ 116°11′37.3″, Xiangshan, Beijing, collected in September 2015), *M. v. melanantha* parasitizing *S. melanantha* (#1296; 27°01′50.6″ 100°10′41.6″ First Peak, Lijiang, date collected in July 2014), and *M. coronariae* parasitizing *L. flos cuculi* (#1247; GPS 55.7 −4.9, Great Cumbrae Island, UK, collected in 2000). Additionally, we sequenced the a_1_ genome of *M. intermedium* parasitizing *Salvia pratensis* (#1389, GPS 44.3353 N 7.13637 E, Cuneo, Italy, collected in July 2011). Samples were collected before the publication of laws regarding the Nagoya protocol in the countries of collection. DNA extraction and sequencing based on Pacific Bioscience long-read sequencing was performed as described previously (Branco, et al. 2017; Branco, et al. 2018).Sequencing was carried out at UCSD IGM Genomics Center (San Diego, USA) with the P6 C4 chemistry. Coverage was between 24x and 35x for all genomes, except for *M. intermedium* for which it was 199x.

### Assembly of new genomes and assembly improvement for *M. scabiosae* infecting *Knautia arvensis*

We converted the bax.h5 files from each smart cell into one bam file using bax2bam with the “--subread” option (https://github.com/PacificBiosciences/bax2bam), and all bam files from the same sequencing genome into one fastq file using bam2fastx (https://github.com/PacificBiosciences/bam2fastx). We generated the assembly using canu (Koren, et al. 2017) with the parameters “genomeSize=30m” and “-pacbio-raw”. We used pbalign (version 0.3.0) with the blasr algorithm (Chaisson and Tesler 2012) to realign the raw reads onto the assembly (indexed with samtools faidx (Li, et al. 2009) and then used the output bam file into quiver (Chin, et al. 2013) to polish the assembly basecalling (https://github.com/PacificBiosciences/GenomicConsensus). Default parameters were used when not detailed in the text (see the Table S1 for further assembly statistics).

The previous genome assemblies of the a_1_ and a_2_ genome of *M. scabiosae* were based on five PacBio movies (Branco, et al. 2018). We re-assembled the genome using three additional movies, from the same strain, and generated using the same PacBio technology for a total coverage of 506x for the a_1_ genome and 803x for the a_2_ genome. The assemblies were substantially improved, as the contig numbers were reduced from 123 and 147 to 26 and 20 for the a_1_ and a_2_ mating type, respectively (see the Table S2 for further assembly statistics).

### Gene models, orthologous group reconstruction, transposable elements, centromeres

As for the previously published *Microbotryum* genomes (Badouin, et al. 2015; Branco, et al. 2017; Branco, et al. 2018), the protein-coding gene models were predicted with EuGene (version 4.2a;(Foissac, et al. 2008), trained for M*icrobotryum*. Similarities to the fungal subset of the uniprot database (The UniProt 2011) plus the *M. lychnidis-dioicae* Lamole proteome (Badouin, et al. 2015)were integrated into EuGene for the prediction of gene models. We obtained the orthologous groups based on all-vs-all blastp high scoring pairs parsed with orthAgogue -u option (Ekseth, et al. 2014) and clustered with mcl (Dongen 2000) setting to 1.5 the inflation parameter.Transposable elements were obtained from a previous study (Hartmann, et al. 2018).Centromeres were identified as regions with the highest density of the centromeric repeats described in *M. lychnidis-dioicae* (Badouin, et al. 2015). In the most fragmented assemblies (e.g. *M. coronariae*), the centromeres on the mating-type chromosomes could not be localized, but the centromere localization was only used in the *M. lagerheimii* assembly to assess whether chromosome fissions occurred at centromeres.

### Evolutionary scenarios of the mating-type chromosomes

For the figures S1-S3, we represented genomic data using Circos (Krzywinski 2009) with the following tracks: (i) contig, (ii) dS=0 (indicating recombination in these selfing species as selfing leads to homozygosity), (iii) all genes, (iv) transposable elements, centromeric repeats.Both the d_S_=0 and all gene tracks were filtered for TEs and centromeric repeats, using their merged coordinates with bedtools (Quinlan and Hall 2010). Homozygosity (d_S_=0) was used to infer recombining regions on mating-type chromosomes following previous studies (Branco, et al. 2017; Branco, et al. 2018). The comparisons of mating-type chromosomes of the species with linked HD and PR loci to the ancestral state with HD and PR loci on different chromosomes (using the *M. intermedium* genome as a proxy for the ancestral state) allowed us to infer the chromosomal rearrangements having linked HD and PR loci, as previously done (Branco, et al. 2017; Branco, et al. 2018). Non-recombining regions were identified as genomic regions with non-zero levels of differentiation between mating-type chromosomes as previously done (Branco, et al. 2017; Branco, et al. 2018).

### RNAseq experiment and expression level quantification

We generated RNAseq data for *M. intermedium* under two different conditions: i) haploid cells of a single mating type grown on rich medium and then mixed in equal proportions of the two mating types before RNA extraction (“H” condition), and ii) mixtures of cells of the two mating types under mating conditions (“M” condition). Total RNA was isolated from haploid cells using the Qiagen RNeasy Mini Kit. For the rich medium condition, haploid strains (a_1_ or a_2_) were streaked on Potato Dextrose Agar (PDA) and grown for 48 hours at 22°C. Cells were scraped, ground in liquid nitrogen, and total RNA extracted following the manufacturer’s protocol. For RNAseq analysis, equal amounts of total RNA individually isolated from a_1_ and a_2_ cultures were pooled. For the mating condition, haploid a_1_ and a_2_ cultures were first grown separately in yeast extract peptone dextrose broth overnight. Cell density was measured with a spectrophotometer. The concentration of each culture was adjusted to O.D._600_ 1.0. Equal volumes of the two haploid cultures were mixed and plated on water agar. These plates were incubated for four days at 14°C, after which wet mounts were prepared from each mated plate to verify the presence of conjugation tubes, indicating active mating behaviour. Cells were scraped. Total RNA was isolated from haploid cells using the Qiagen RNeasy Mini Kit. After total RNA isolation, several quality control measures were taken. Concentration and purity were assessed using a NanoDrop 2000 spectrophotometer (Thermo Scientific); 260/280 and 260/230 ratios greater than 1.8 were considered satisfactory for RNAseq application. Additionally, cDNA was prepared and used as template for intron-spanning primers in PCR reactions to verify the lack of genomic DNA contamination. Bioanalyzer analysis was completed to detect intact 18S and 23S RNA as a measure of overall RNA quality. After passing all quality control measurements, RNA was sent for RNAseq analysis to CD Genomics (Shirley, New York). Three replicates were sequenced per condition and means across the three replicates were analysed.

For *M. lychnidis-dioicae*, we used RNAseq data published previously (Perlin, et al. 2015), using only the expression data from the two conditions that corresponded to those analysed in *M. intermedium*, i.e. culture on PDA medium and under mating conditions. List and statistics of the RNAseq experiments used in this study are provided in Table S4. We controlled the quality of the RNAseq experiment data using fastQC (https://github.com/s-andrews/FastQC). We trimmed the sequences using trimmomatic (Bolger et al. 2014); parameters: ILLUMINACLIP:TruSeq3-PE.fa:2:30:10:2:keepBothReads LEADING:3 TRAILING:3 MINLEN:36). We only considered the resulting trimmed paired-end reads for further analysis (Table S3). For each RNAseq dataset, we pseudo-aligned the reads onto the corresponding references and quantified the gene expression levels using Kallisto using the “kallisto quant” command with the “--bias” option (Bray, et al. 2016). For the diploid RNAseq experiment (Table S4), we used the concatenated coding sequence set from a_1_ and a_2_ haploid genomes as reference. For haploid RNAseq experiment, we considered the haploid coding sequence set from the corresponding mating type. For each RNAseq experiment, we removed all the genes with an expression level above the 95^th^ percentile of the expression level distribution to avoid spurious high expression level.

We tested for change in expression level for genes in recombining regions between two distantly-related *Microbotryum* species. We performed Pearson’s correlation tests between *M. intermedium* and *M. lychnidis-dioicae* orthologous genes for the two of RNA-seq experimental replicates that were similar between the two species (*i.e*. cells grown on rich media or mated on water agar (Perlin, et al. 2015); Table S8).

### Identifying optimal codons in *Microbotryum* fungi

We generated a codon usage table for each of the a_1_ *M. lychnidis-dioicae*, a_2_ *M. lychnidis-dioicae*, a_1_ *M. intermedium* and a_2_ *M. intermedium* haploid genomes (scripts available in the public repository) and filtered the genes that overlapped transposable elements that had been previously identified (Hartmann, et al. 2018). We then performed a within-group correspondence analysis (WCA), following the procedure available at http://pbil.univ-lyon1.fr/datasets/charif04/ (Charif, et al. 2005) on each codon usage table. Correspondence analysis is a multivariate statistical method that summarizes codon count data by reducing them to a limited number of variables. WCA more specifically dissociates the effects of different amino-acid compositions from the effects directly related to synonymous codon usage (Charif, et al. 2005). This method adjusts the value for each codon by the average value of all the codons encoding for the same amino acid. We only considered the two first principal components which explained between 16.30% and 16.50% of the variance for the first principal component and between 7.75% and 8.51% for the second principal component (Figure S4). The coordinates of the genes projected onto the first (PC1) and second (PC2) axes were significantly correlated with gene expression level (Pearson's correlation tests; Table S5). We considered the first axis to better characterized the gene expression level variation because the correlations between PC1 gene coordinates and gene expression level were higher than those calculated between PC2 gene coordinates and gene expression level (see correlation’s coefficients in Table S5).

We performed chi-square tests to compare, per amino acid, the codon counts of the 10% of genes having the lowest PC1 coordinates to the codon counts of the 10% of genes having the highest PC1 coordinates, representing the genes with highest and lowest expression, respectively (Figure S3). When the chi-square test was significant (*p* < 0.05), we considered for each amino acid the optimal codon to be the one with the highest residual from the chi-square test (Table S6). We did not consider the ATG (methionine) and TGG (tryptophan) codons as they have no synonymous codons.

### Frequency of optimal codons and GC content statistics

For each coding sequence (i.e., exons) with base pair numbers corresponding to multiple of three, we calculated the frequency of optimal codons (FOP), the GC content of the third position (GC_3_) of each codon and the GC content of the introns of each gene (GC_introns_). We removed from this gene set any gene overlapping with a transposable element.

### Non-synonymous and synonymous divergence estimation

For each pair of allele from alternative mating type in each species, we aligned the protein sequences with muscle v3.8.31 (Edgar 2004) and obtained the codon-based CDS alignments with TranslatorX; d_N_ and d_S_ values were computed in the yn00 program for each gene in each species (Yang and Nielsen 2000). To estimate d_N_/d_S_ values per stratum, we summed the d_N_ and d_S_ values of all genes within each stratum, weighted by gene size, and computed the ratio of these sums.

### Absolute dating of recombination suppression events

We estimated the species and genome compartment divergence times under a calibrated Yule model and a HKY substitution model (kappa=2, estimated frequencies) with 10,000,000 or 20,000,000 mcmc generations in beast v2.5.0 (Bouckaert, et al. 2019). We used three unlinked site models (one per codon position). Clock model and tree parameters were kept linked. We used the split of *M. lychnidis-dioicae* and *M. silenes-dioicae* as a calibration point, with a normal prior with a mean 0.42 MYA and sigma 0.04 (Gladieux, et al. 2011). Beast input files are available in the public repository. Concatenated alignments were obtained from codon-based alignments produced by macse v2.03 -prog alignSequences (Ranwez, et al. 2018) protocol on fully conserved single copy genes. We reconstructed the species and genome compartment maximum likelihood trees with IQTree v2.0.4 (1000 rapid bootstraps and automatic model selection (Minh, et al. 2020)).

For the species divergence we used 3,955 fully conserved single copy autosomal genes (2,732,071 aligned codons). For estimating the dates of recombination suppression, we estimated the divergence time at the genes linked to mating-type loci between alleles associated with a_1_ and a_2_. mating types in ML trees. We used genes ancestrally located on the long arm of the HD chromosome (corresponding to the white stratum, with 179 genes and 129,122 aligned codons, and the light brown stratum, with 92 genes and 78,627 aligned codons), the region between the HD loci and the centromere (8 genes and 23,115 aligned codons), the region between the HD loci and the short arm telomere (12 genes and 9,165 aligned codons), the short arm of PR chromosome (11 genes and 11,865 aligned codons), the region between the PR loci and the centromere (23 genes and 19,944 aligned codons) and the region extending from the PR locus towards the long arm telomere (shared gene set between the PR associated non-recombining region of *M. silenes-acaulis* and the red, light blue and pink strata, with 14 genes and 9,952 aligned codons).

### Statistics

We tested, for each of the 26 *Microbotryum* genomes (one genome per mating type for 13 species), whether optimal codons were less common in the genes in distinct genomic compartments (PARs and the different evolutionary strata with different ages of recombination suppression) than in autosomal genes. To control for an effect of GC-biased gene conversion on differences in GC level among genome compartments, we used the difference in GC content between coding and intron sequences across genes as a covariate in an ANCOVA that considered the status of codons (optimal versus non-optimal) as the response variable, genomic compartment as classification variable and the difference in GC content between coding and intron sequences across genes as the covariate. The interaction effect tested for difference in slopes of the relationship between codon usage and the covariate among genome compartments. Using the glm function in R, we gave as input the frequency of optimal codons weighted by the total number of codons per gene (using the “weight” option in the glm function). After performing a logistic regression assuming binomial distribution and a logit link, we noticed that predicted frequencies of optimal codons were all lower than the raw frequency of optimal codons which could indicate overdispersion issues. We therefore performed the logistic regression assuming a quasibinomial distribution, the only difference being in the estimation of the dispersion parameter to correct, among other things, the p-values. The estimated dispersion parameter was high for the logistic regression model of each genome (Ф > 2), so we choose the quasibinomial distribution to interpret the data. Choosing a quasibinomial distribution did not, however, change the log(odds) estimated by the model.

To assess how the frequency of optimal codons, the ratio of non-synonymous over synonymous mutations (d_N_/d_S_) and the number of non-synonymous mutations (d_N_) varied over time since recombination suppression in non-recombining regions, we performed regressions using R. We used species as blocks and, for the optimal codon analysis, ancestral gene expression level as a covariable. We compared linear, logarithmic or quadratic regressions based on Akaike information criterion (AIC). For d_N_/d_S_ and d_N_ analyses, we analysed only non-recombining regions, i.e. we did not include autosomes or pseudo-autosomal regions, as nearly no synonymous or non-synonymous mutations were observed in these genomic compartments, the species being highly homozygous. Because the status for a codon as optimal or not is a binomial variable, we performed logistic regressions for this trait using generalized linear models.

To visualise the variation in the frequency of optimal codons or d_N_/d_S_ for all strata, including those for which we had no solid age estimates because of their small sizes, we plotted the values as boxplots and as a function of d_S_.

## Author contribution

Conception and design, TG, MEH, FC; Formal analyses, FC, RCRV, PJ, MD; Interpretation, FC, TG, MEH, RCRV, JAS; Data acquisition, MHP, RMW, FC, TG, RCRV; Original draft, FC, TG; Final draft, FC, TG, MEH, RCRV, JAS, PJ; Revision, FC, TG, RCRV, MEH, MHP, RMW. All authors read and approved the final version.

## Acknowledgements

We thank Hui Tang and Janis Antonovics for help with sample collection. We thank Maxime Dubart and Sylvain Billiard for great help and discussion about statistical analyses. This work was supported by the National Institute of Health (NIH) grant number R15GM119092 to M. E. H., and the Louis D. Foundation award and EvolSexChrom ERC advanced grant number 832352 to T. G. We thank Hector Mendoza for help with RNA isolation. We thank Kurt Hasselman, John Bain and Hui Tang who collected the strains (involving a confrontation with a grizzly bear for John Bain). Support for R.M.W. and RNA isolation was additionally from NIH [sub-award #OGMB131493D1] to M.H.P. from [P20GM103436] to [Nigel Cooper, PI] and also NSF/IRES Award (1824851) to M.H.P. The contents of this work are solely the responsibility of the authors and do not represent the official views of the NIH.

## Additional files

1-All supplementary Tables as a single xls file (different Tables as sheets)

2-All supplementary Figures as a single PDF file

## Supplemental Figure legends

**Figure S1: Synteny between mating-type chromosomes of alternative mating types for the *Microbotryum* species A) *M. lychnis flos-cuculi*, B) *M. v. melanantha* and C) *M. v. tatarinowii*.** For each species, we show synteny plots between a_1_ and a_2_ mating-type chromosomes (top left), synteny plots between a_1_ mating-type chromosome and a_1_ mating-type chromosome of *M. intermedium* (middle panel) and synteny plots between the a_2_ mating-type chromosome of the focal species and the a_2_ mating-type chromosome of *M. intermedium* (right panel). On the synteny plots on the left, the five tracks represent, from the inner most to the outermost: i) centromeric repeats as red lines, ii) genes as grey lines, iii) transposable elements as green lines, iv) genes with zero synonymous substitutions between mating types as black lines, v) contigs coloured according to species or for *M. intermedium* in blue for the HD chromosome, purple for the PR chromosome, grey for the autosomes, with yellow regions indicating centromere-specific repeats, and with the axis indicating megabases. The purple circle indicates the PR locus and the blue circle the HD locus. Fine blue and orange lines link regions with collinearity extending over 2 kb, the latter corresponding to inversions. Areas without links correspond to highly rearranged regions. On the synteny plots at right, only the tracks i to iii are represented. The small contigs within the non-recombining region in some species could not be oriented, but this does not affect our inferences of chromosomal rearrangements or degeneration measures.

**Figure S2: Synteny and allelic divergence analysis between mating-type chromosomes of alternative mating types for the species *Microbotryum intermedium*.** We show synteny plots between a_1_ and a_2_ PR mating-type chromosome (left top panel), synteny plots between a_1_ and a_2_ HD mating-type chromosomes (right top panel), and the synonymous divergence calculated between alternative mating types and plotted along the *M. intermedium* PR and HD mating-type chromosome gene order (bottom left and right panel, resp.). Zero d_S_ between mating-type chromosomes indicate recombination in these selfing species and the level of d_S_ between mating-type chromosomes increases with time since recombination suppression. Different evolutionary strata, i.e., different steps extending recombination suppression, are indicated by different colours. Divergence between the a_1_ and a_2_ pheromone receptor (PR) genes was too extensive, and d_S_ could not be calculated (depicted as “Un” for unalignable). On the synteny plots, the five tracks represent, from the inner most to the outermost: i) centromeric repeats as brown lines, ii) genes as black lines, iii) transposable elements as green lines, iv) genes with zero synonymous substitutions between mating types as black lines, v) contigs coloured in blue for the HD chromosome and purple for the PR chromosome, with the axis indicating megabases. The purple circle indicates the PR locus and the blue circle the HD locus. Fine blue and orange lines link regions with collinearity extending over 2 kb, the latter corresponding to inversions.

**Figure S3**: **Synteny and allelic divergence analysis between mating-type chromosomes of alternative mating types for the species *M. scabiosae* (A) and *Microbotryum v. caroliniana* (B).** For each species, we show synteny plots between a_1_ and a_2_ mating-type chromosomes (top panel), synteny plots between a_1_ HD and PR mating-type chromosomes of *M. intermedium* and a_1_ or a_2_ mating-type chromosomes of the focal species (right and left middle panels, respectively), and the synonymous divergence calculated between alternative mating types and plotted along the *M. intermedium* mating-type chromosome gene order (bottom panel). Zero d_S_ between mating-type chromosomes indicate recombination in these selfing species and the level of d_S_ between mating-type chromosomes increases with time since recombination suppression. On the top left figure in B), the light blue evolutionary stratum is indicated by a light blue box on the outer track. On the synteny plots at left, the five tracks represent, from the inner most to the outermost: i) centromeric repeats as brown lines, ii) genes as black lines, iii) transposable elements as green lines, iv) genes with zero synonymous substitutions between mating types as black lines, v) contigs colored according to species or for *M. intermedium* in blue for the HD chromosome, purple for the PR chromosome, grey for the autosomes, with yellow regions indicating centromere-specific repeats, and with the axis indicating megabases. The purple circle indicates the PR locus and the blue circle the HD locus. Fine blue and orange lines link regions with collinearity extending over 2 kb, the latter corresponding to inversions. Areas without links correspond to highly rearranged regions. On the synteny plots at right, only the tracks i to iii are represented.

**Figure S4**: Within-correspondence analysis plots. A) WCA plots for the a_1_ and a_2_ genomes of *Microbotryum intermedium* (resp. left and right top panel) and a_1_ and a_2_ genomes of *M. lychnidis-dioicae* (resp. left and right bottom panel). B) Percentage of variance explained for each correspondence analysis axis for the a_1_ and a_2_ genomes of *M. intermedium* (resp. left and right top panel) and a_1_ and a_2_ genomes of *M. lychnidis-dioicae* (resp. left and right bottom panel).

**Figure S5: Data distribution across genomic compartments in *Microbotryum*.** A) Distribution of the frequency of optimal codons. B) Distribution of the difference in GC content between coding sequences and introns. C) Distribution of the expression level of *M. intermedium* orthologous genes. The gene expression level distribution was truncated at the 90^th^ percentile for visualisation purpose and was not represented for the genomes a_1_ and a_2_ *M. intermedium*. Boxplot colours refer to the genomic compartment.

**Figure S6**: **Analysis of the frequency of optimal codons in all *Microbotryum* genomes using the genomic compartments considered in the main text.** A) Distribution of the frequency of optimal codons across genomic compartments. B) Distribution of the predicted frequency of optimal codons predicted by the logistic regression. In A) and B), boxplot colours refer to the genomic compartment. For each genomic compartment, the red dot indicates the mean frequency of optimal codons, the sample size (N) is labelled on top of the corresponding boxplot, as well as the significance level of the difference between a given genomic compartment and the autosomes (NS: non-significant; “.”: p < 0.1; *: p < 0.05; **: p-value < 0.01; ***: p < 0.001). C) Frequency of optimal codons predicted by the logistic regression along the difference of GC content between coding and intronic sequences. Significant differences in slope between a genomic compartment and the autosomes are indicated by a solid line and a filled dot, while non-significant differences in slope are indicated by a dotted line and an open dot.

**Figure S7: Analysis of the frequency of optimal codons in all *Microbotryum* genomes using all the genomic compartments.** A) Distribution of the frequency of optimal codons across genomic compartments. B) Distribution of the predicted frequency of optimal codons predicted by the logistic regression. In A) and B), boxplot colours refer to the genomic compartment. For each genomic compartment, the red dot indicates the mean frequency of optimal codons, the sample size (N) is labelled on top of the corresponding boxplot, as well as the significance level of the difference between a given genomic compartment and the autosomes (NS: non-significant; “.”: p-value < 0.1; *: p-value < 0.05; **: p-value < 0.01; ***: p-value < 0.001). C) Frequency of optimal codons predicted by the logistic regression as a function of the difference of GC content between coding and intronic sequences. Significant differences in slope between a genomic compartment and the autosomes are indicated by a solid line and a filled dot, while non-significant differences in slope are indicated by a dotted line and an open dot.

**Figure S8: Tempo of degeneration for all genomic compartments and species, as a function of synonymous divergence (d_S_) between mating-type chromosomes in the focal genomic compartments.** Degeneration is plotted in terms of (A) frequencies of optimal codons, (B) non-synonymous differences over synonymous ratio (d_N_/d_S_) between alleles of alternative mating-type chromosomes, (C) non-synonymous differences (d_N_) between alleles of alternative mating-type chromosomes.

**Figure S9: Frequency of optimal codons for all genes in all *Microbotryum* genomes as a function of the expression level in *M. intermedium* as a proxy for the ancestral expression level.** The color indicates the age of the recombination suppression event of the genomic compartment in which the focal gene resides. The *M. intermedium* expression level used is from the H condition (haploid cells of a single mating type on rich medium) in A and from the M condition (mixtures of cells of the two mating types under mating conditions) in B.

## Cited literature

Bachtrog D. 2005. Sex chromosome evolution: Molecular aspects of Y-chromosome degeneration in Drosophila. Genome Research 15: 1393–1401. doi: 10.1101/gr.3543605

Bachtrog D. 2008. The temporal dynamics of processes underlying Y chromosome degeneration. Genetics 179: 1513–1525. doi: 10.1534/genetics.107.084012

Bachtrog D. 2013. Y-chromosome evolution : emerging insights into processes of Y-chromosome degeneration. Nature Reviews Genetics 14: 113–124. doi: 10.1038/nrg3366

Badouin H, Hood ME, Gouzy J, Aguileta G, Siguenza S, Perlin MH, Cuomo CA, Fairhead C, Branca A, Giraud T. 2015. Chaos of rearrangements in the mating-type chromosomes of the anther-smut fungus *Microbotryum lychnidis-dioicae*. Genetics 200: 1275–1284. doi: 10.1534/genetics.115.177709

Bartolomé C, Charlesworth B. 2006. Evolution of amino-acid sequences and codon usage on the *Drosophila miranda* neo-sex chromosomes. Genetics 174: 2033–2044. doi: 10.1534/genetics.106.064113

Bellott DW, Hughes JF, Skaletsky H, Brown LG, Pyntikova T, Cho TJ, Koutseva N, Zaghlul S, Graves T, Rock S, Kremitzki C, Fulton RS, Dugan S, Ding Y, Morton D, Khan Z, Lewis L, Buhay C, Wang Q, Watt J, Holder M, Lee S, Nazareth L, Rozen S, Muzny DM, Warren WC, Gibbs RA, Wilson RK, Page DC. 2014. Mammalian y chromosomes retain widely expressed dosage-sensitive regulators. Nature 508: 494–499. doi: 10.1038/nature13206

Bergero R, Charlesworth D. 2011. Preservation of the Y transcriptome in a 10-Million-year-old plant sex chromosome system. Current Biology 21: 1470–1474.

Berlin S, Ellegren H. 2006. Fast accumulation of nonsynonymous mutations on the female-specific W chromosome in birds. J. Mol. Evol. 62: 66–72.

Bernstein H, Byers GS, Michod RE 1981. Evolution of sexual reproduction: importance of DNA repair, complementation, and variation. Am. Nat. 117: 537–549.

Betancourt AJ, Welch JJ. 2009. Report reduced effectiveness of selection caused by a lack of recombination. Current Biology 19: 655–660. doi: 10.1016/j.cub.2009.02.039

Bianchi NO. 2009. Y chromosome structural and functional changes in human malignant diseases. Mutation research 682: 21–27. doi: 10.1016/j.mrrev.2009.02.001

Bouckaert R, Vaughan TG, Barido-Sottani J, Duchêne S, Fourment M, Gavryushkina A, Heled J, Jones G, Kühnert D, De Maio N, Matschiner M, Mendes FK, Müller NF, Ogilvie HA, du Plessis L, Popinga A, Rambaut A, Rasmussen D, Siveroni I, Suchard MA, Wu C-H, Xie D, Zhang C, Stadler T, Drummond AJ. 2019. BEAST 2.5: An advanced software platform for Bayesian evolutionary analysis. PLOS Computational Biology 15: 1–28. doi: 10.1371/journal.pcbi.1006650

Branco S, Badouin H, Rodríguez de la Vega RC, Gouzy J, Carpentier F, Aguileta G, Siguenza S, Brandenburg J-T, Coelho MA, Hood ME, Giraud T. 2017. Evolutionary strata on young mating-type chromosomes despite the lack of sexual antagonism. Proceedings of the National Academy of Sciences 114: 7067–7072. doi: 10.1073/pnas.1701658114

Branco S, Carpentier F, Rodríguez de la Vega RC, Badouin H, Snirc A, Le Prieur S, Coelho MA, de Vienne DM, Hartmann FE, Begerow D, Hood ME, Giraud T. 2018. Multiple convergent supergene evolution events in mating-type chromosomes. Nature Communications 9: 2000. doi: 10.1038/s41467-018-04380-9

Bray NL, Pimentel H, Melsted P, Pachter L. 2016. Near-optimal probabilistic RNA-seq quantification. Nature Biotechnology 34: 525–527. doi: 10.1038/nbt.3519

Brion C, Caradec C, Pflieger D, Friedrich A, Schacherer J. 2020. Pervasive phenotypic impact of a large non-recombining introgressed region in yeast. Molecular Biology and Evolution 37: 2520–2530.

Carlini DB. 2004. Experimental reduction of codon bias in the *Drosophila* alcohol dehydrogenase gene results in decreased ethanol tolerance of adult flies. Journal of Evolutionary Biology 17: 779–785. doi: 10.1111/j.1420-9101.2004.00725.x

Carpentier F, Rodríguez RC, Vega D, Branco S, Snirc A, Coelho MA, Hood ME, Giraud T, Evolution ES, Paris-saclay U, Ciências DD, Ciências FD, Lisboa UND. 2019. Convergent recombination cessation between mating-type genes and centromeres in selfing anther-smut fungi. Genome Research 29: 944–953. doi: 10.1101/gr.242578.118.5

Chaisson MJ, Tesler G. 2012. Mapping single molecule sequencing reads using basic local alignment with successive refinement (BLASR): application and theory. BMC Bioinformatics 13: 238.

Charif D, Thioulouse J, Lobry JR, Perrière G. 2005. Online synonymous codon usage analyses with the ade4 and seqinR packages. Bioinformatics 21: 545–547. doi: 10.1093/bioinformatics/bti037

Charlesworth B, Charlesworth D. 2000. The degeneration of Y chromosomes. Philosophical Transactions of the Royal Society B-Biological Sciences 355: 1563–1572. doi: 10.1098/rstb.2000.0717

Charlesworth B, Charlesworth D 1978. A model for the evolution of dioecy and gynodioecy. The American Naturalist 112: 975–997. doi: 10.1017/CBO9781107415324.004

Charlesworth D. 2021. The timing of genetic degeneration of sex chromosomes. Phil. Trans. R. Soc. B 376: 20200093.

Chibalina MV, Filatov DA. 2011. Plant y chromosome degeneration is retarded by haploid purifying selection. Current Biology 21: 1475–1479. doi: 10.1016/j.cub.2011.07.045

Chin CS, Alexander DH, Marks P, Klammer AA, Drake J, Heiner C, Clum A, Copeland A, Huddleston J, Eichler EE, Turner SW, Korlach J. 2013. Nonhybrid, finished microbial genome assemblies from long-read SMRT sequencing data. Nature Methods 10: 536.

Coelho MA, Bakkeren G, Sun S, Hood ME, Giraud T. 2017. Fungal Sex: The Basidiomycota. Microbiology Spectrum 5: 1–30. doi: 10.1128/microbiolspec.FUNK-0046-2016

Corcoran P, Anderson JL, Jacobson DJ, Sun Y, Ni P, Lascoux M, Johannesson H. 2016. Introgression maintains the genetic integrity of the mating-type determining chromosome of the fungus *Neurospora tetrasperma*. Genome Research 26: 486–498. doi: 10.1101/gr.197244.115

Dongen SV. 2000. A cluster algorithm for graphs. Information Systems [INS]: 1–40. doi: INS-R0010

Duhamel M, Carpentier F, Begerow D, Hood ME, Rodríguez de la Vega RC, Giraud T. 2022. Onset and stepwise extensions of recombination suppression are common in mating-type chromosomes of Microbotryum anther-smut fungi. J Evol Biol BioRxiv DOI: 10.1101/2021.10.29.466223

Duret L. 2002. Evolution of synonymous codon usage in metazoans. Current Opinion in Genetics and Development 12: 640–649. doi: 10.1016/S0959-437X(02)00353-2

Duret L, Galtier N. 2009. Biased gene conversion and the evolution of mammalian genomic landscapes. Annual Review of Genomics and Human Genetics 10: 285–311. doi: 10.1146/annurev-genom-082908-150001

Edgar RC. 2004. MUSCLE: Multiple sequence alignment with high accuracy and high throughput. Nucleic Acids Research 32: 1792–1797. doi: 10.1093/nar/gkh340

Ekseth OK, Kuiper M, Mironov V. 2014. OrthAgogue: an agile tool for the rapid prediction of orthology relations. Bioinformatics 30: 734–736. doi: 10.1093/bioinformatics/btt582

Fisher RA. 1930. The genetical theory of natural selection.

Foissac S, Gouzy J, Rombauts S, Mathe C, Amselem J, Sterck L, de Peer VY, Rouze P, Schiex T. 2008. Genome annotation in plants and fungi: EuGene as a model platform. Current Bioinformatics 3: 87–97.

Fontanillas E, Hood ME, Badouin H, Petit E, Barbe V, Gouzy J, De Vienne DM, Aguileta G, Poulain J, Wincker P, Chen Z, Toh SS, Cuomo CA, Perlin MH, Gladieux P, Giraud T. 2015. Degeneration of the nonrecombining regions in the mating-type chromosomes of the anther-smut fungi. Molecular Biology and Evolution 32: 928–943. doi: 10.1093/molbev/msu396

Gladieux P, Vercken E, Fontaine MC, Hood ME, Jonot O, Couloux A, Giraud T. 2011. Maintenance of fungal pathogen species that are specialized to different hosts: allopatric divergence and introgression through secondary contact. Molecular Biology and Evolution 28: 459–471. doi: 10.1093/molbev/msq235

Gutiérrez-Valencia J, Hughes PW, Berdan EL, Slotte T. 2021. The genomic architecture and evolutionary fates of supergenes. Genome Biology and Evolution 13. doi: 10.1093/gbe/evab057

Hartmann FE, Duhamel M, Carpentier F, Hood ME, Foulongne-Oriol M, Silar P, Malagnac F, Grognet P, Giraud T. 2021. Recombination suppression and evolutionary strata around mating-type loci in fungi: documenting patterns and understanding evolutionary and mechanistic causes. New Phytologist 229: 2470–2491. doi: 10.1111/nph.17039

Hartmann FE, Rodríguez de la Vega RC, Brandenburg J-T, Carpentier F, Giraud T. 2018. Gene presence–absence polymorphism in castrating anther-smut fungi: recent gene gains and phylogeographic structure. Genome Biology and Evolution 10: 1298–1314. doi: 10.1093/gbe/evy089

Hartmann FE, Rodríguez de la Vega RC, Gladieux P, Ma W-J, Hood ME, Giraud T. 2020. Higher gene flow in sex-related chromosomes than in autosomes during fungal divergence. Molecular Biology and Evolution 37: 668–682. doi: 10.1093/molbev/msz252

Hill WG, Robertson A 1966. The effect of linkage on limits to artificial selection. Genetics Research 8: 269–294. doi: 10.1017/S001667230800949X

Hollister JD, Gaut BS. 2009. Epigenetic silencing of transposable elements: A trade-off between reduced transposition and deleterious effects on neighboring gene expression. Genome Research 19: 1419–1428. doi: 10.1101/gr.091678.109

Hood ME, Antonovics J. 2000. Intratetrad mating, heterozygosity, and the maintenance of deleterious alleles in Microbotryum violaceum (= Ustilago violacea). Heredity 85: 231–241. doi: 10.1046/j.1365-2540.2000.00748.x

Hood ME, Mena-Alí JI, Gibson AK, Oxelman B, Giraud T, Yockteng R, Arroyo MTK, Conti F, Pedersen AB, Gladieux P, Antonovics J. 2010. Distribution of the anther-smut pathogen *n* on species of the Caryophyllaceae. New Phytologist 187: 217–229. doi: 10.1111/j.1469-8137.2010.03268.x

Hough J, Hollister JD, Wang W, Barrett SCH, Wright SI. 2014. Genetic degeneration of old and young y chromosomes in the flowering plant Rumex hastatulus. Proceedings of the National Academy of Sciences of the United States of America 111: 7713–7718. doi: 10.1073/pnas.1319227111

Hughes JF, Rozen S. 2012. Genomics and genetics of human and primate Y chromosomes. Annual Review of Genomics and Human Genetics 13: 83–108. doi: 10.1146/annurev-genom-090711-163855

Ikemura T 1981. Correlation between the abundance of *Escherichia coli* transfer RNAs and the occurrence of the respective codons in its protein genes: A proposal for a synonymous codon choice that is optimal for the *E. coli* translational system. Journal of Molecular Biology 151: 389–409. doi: 10.1016/0022-2836(81)90003-6

Jay P, Chouteau M, Whibley A, Bastide H, Parrinello H, Llaurens V, Joron M. 2021a. Mutation load at a mimicry supergene sheds new light on the evolution of inversion polymorphisms. Nature Genetics 53: 288–293. doi: 10.1038/s41588-020-00771-1

Jay P, Tezenas E, Giraud T. 2021b. A deleterious mutation-sheltering theory for the evolution of sex chromosomes and supergenes. bioRxiv: 2021.2005.2017.444504. doi: 10.1101/2021.05.17.444504

Kliman RM, Hey J 1993. Reduced natural selection associated with low recombination in *Drosophila melanogaster*. Mol. Biol. Evol. 10: 1239–1258.

Koren S, Walenz BP, Berlin K, Miller JR, Bergman NH, Phillippy AM. 2017. Canu: scalable and accurate long-read assembly via adaptive k-mer weighting and repeat separation. Genome Research 27: 722–736.

Kostka D, Hubisz MJ, Siepel A, Pollard KS. 2012. The role of GC-biased gene conversion in shaping the fastest evolving regions of the human genome. Molecular Biology and Evolution 29: 1047–1057. doi: 10.1093/molbev/msr279

Krasovec M, Chester M, Ridout K, Filatov DA. 2018. The mutation rate and the age of the sex chromosomes in *Silene latifolia*. Current Biology 28: 1832–1838.e1834. doi: 10.1016/j.cub.2018.04.069

Krzywinski M. 2009. Circos: an information aesthetic for comparative genomics. Genome Res 19: 1639–1645. doi: 10.1101/gr.092759.109.19

Lahn BT, Page DC 1999. Four evolutionary strata on the human X chromosome. Science (New York, N.Y.) 286: 964–967. doi: 10.1126/science.286.5441.964

Lampson BL, Pershing NLK, Prinz JA, Lacsina JR, Marzluff WF, Nicchitta CV, MacAlpine DM, Counter CM. 2013. Rare codons regulate KRas oncogenesis. Current Biology 23: 70–75. doi: 10.1016/j.cub.2012.11.031

Lassalle F, Périan S, Bataillon T, Nesme X. 2015. GC-content evolution in bacterial genomes : the biased gene conversion hypothesis expands. PLoS Genet 11: e1004941. doi: 10.1371/journal.pgen.1004941

Lee Y, Kim C, Park Y, Pyun J-A, Kwack K. 2016. Next generation sequencing identifies abnormal Y chromosome and candidate causal variants in premature ovarian failure patients. Genomics 108: 209–215. doi: https://doi.org/10.1016/j.ygeno.2016.10.006

Lenormand T, Roze D. 2021. Y recombination arrest and degeneration in the absence of sexual dimorphism. bioRxiv: 2021.2005.2018.444606. doi: 10.1101/2021.05.18.444606

Li H, Handsaker B, Wysoker A, Fennell T, Ruan J, Homer N, Marth G, Abecasis G, Durbin R, Genome Project Data Processing Subgroup, Li H, Handsaker B, Wysoker A, Fennell T, Ruan J, Homer N, Marth G, Abecasis G, Durbin R. 2009. The Sequence alignment/map (SAM) format and SAMtools. Bioinformatics 25: 2078–2079.

Li SF, Zhang GJ, Yuan JH, Deng CL, Gao WJ. 2016. Repetitive sequences and epigenetic modification: inseparable partners play important roles in the evolution of plant sex chromosomes. Planta 243: 1083–1095. doi: 10.1007/s00425-016-2485-7

Lutz M, Piatek M, Kemler M, Chlebicki A, Oberwinkler F 2008. Anther smuts of Caryophyllaceae: molecular analyses reveal further new species. Mycological research 112: 1280–1296. doi: 10.1016/j.mycres.2008.04.010

Ma W-J, Carpentier F, Giraud T, Hood M 2020. Differential gene expression between fungal mating types is associated with sequence degeneration. Genome Biology and Evolution. doi: 10.1093/gbe/evaa028

Ma W-J, Veltsos P. 2021. The diversity and evolution of sex chromosomes in frogs. Genes 12: 483. doi: 10.3390/genes12040483

Machado HE, Lawrie DS, Petrov DA. 2020. Pervasive strong selection at the level of codon usage bias in *Drosophila melanogaster*. Genetics: genetics.302542.302019. doi: 10.1534/genetics.119.302542

Marais G. 2003. Biased gene conversion: implications for genome and sex evolution. TRENDS in Genetics 19: 330–338. doi: 10.1016/j.aqpro.2013.07.003

Maynard Smith J, Haigh J 1974. The hitch-hiking effect of a favourable gene. Genet. Res. 23: 22–35. doi: 10.1017/S0016672308009579

Minh BQ, Schmidt HA, Chernomor O, Schrempf D, Woodhams MD, von Haeseler A, Lanfear R. 2020. IQ-TREE 2: New models and efficient methods for phylogenetic inference in the genomic era. Molecular Biology and Evolution 37: 1530–1534. doi: 10.1093/molbev/msaa015

Mrackova M, Nicolas M, Hobza R, Negrutiu I, Monéger F, Widmer A, Vyskot B, Janousek B. 2008. Independent origin of sex chromosomes in two species of the genus *Silene*. Genetics 179: 1129–1133. doi: 10.1534/genetics.107.085670

Müller HJ. 1932. Some genetic aspects of sex. The American Naturalist 66: 118–138.

Nadeau NJ. 2016. Genes controlling mimetic colour pattern variation in butterflies. Current Opinion in Insect Science 17: 24–31. doi: 10.1016/j.cois.2016.05.013

Nicolas M, Marais GAB, Hykelova V, Janousek B, Laporte V, Vyskot B, Mouchiroud D, Negrutiu I, Charlesworth D, Monéger F. 2004. A gradual process of recombination restriction in the evolutionary history of the sex chromosomes in dioecious plants. PLoS Biology 3: e4.

Nieuwenhuis BPS, Billiard S, Vuilleumier S, Petit E, Hood ME, Giraud T. 2013. Evolution of uni-and bifactorial sexual compatibility systems in fungi. Heredity 111: 445–455. doi: 10.1038/hdy.2013.67

Novoa EM, Ribas de Pouplana L. 2012. Speeding with control: Codon usage, tRNAs, and ribosomes. TRENDS in Genetics 28: 574–581. doi: 10.1016/j.tig.2012.07.006

Ohno S. 1967. Sex chromosomes and sex-linked genes.

Otto SP, Lenormand T. 2002. Resolving the paradox of sex and recombination. Nature Reviews Genetics 3: 252–261. doi: 10.1038/nrg761

Papadopulos AST, Chester M, Ridout K, Filatov DA. 2015. Rapid Y degeneration and dosage compensation in plant sex chromosomes. Proceedings of the National Academy of Sciences 112: 201508454. doi: 10.1073/pnas.1508454112

Perlin MH, Amselem J, Fontanillas E, Toh SS, Chen Z, Goldberg J, Duplessis S, Henrissat B, Young S, Zeng Q, Aguileta G, Petit E, Badouin H, Andrews J, Razeeq D, Gabaldón T, Quesneville H, Giraud T, Hood ME, Schultz DJ, Cuomo CA. 2015. Sex and parasites: Genomic and transcriptomic analysis of *Microbotryum lychnidis-dioicae*, the biotrophic and plant-castrating anther smut fungus. BMC genomics 16: 1–24. doi: 10.1186/s12864-015-1660-8

Pessia E, Popa A, Mousset S, Rezvoy C, Duret L, Marais GAB. 2012. Evidence for widespread GC-biased gene conversion in eukaryotes. Genome Biol. Evol. 4: 675–682. doi: 10.1093/gbe/evs052

Post LE, Strycharz GD, Nomura M, Lewis H, Dennis PP 1979. Nucleotide sequence of the ribosomal protein gene cluster adjacent to the gene for RNA polymerase subunit β in Escherichia coli. Proceedings of the National Academy of Sciences of the United States of America 76: 1697–1701. doi: 10.1073/pnas.76.4.1697

Qiu S, Bergero R, Zeng K, Charlesworth D. 2011. Patterns of codon usage bias in *Silene latifolia*. Molecular Biology and Evolution 28: 771–780. doi: 10.1093/molbev/msq251

Quinlan AR, Hall IM. 2010. BEDTools: a flexible suite of utilities for comparing genomic features. Bioinformatics 26: 841–842. doi: 10.1093/bioinformatics/btq033

Ranwez V, Douzery EJP, Cambon C, Chantret N, Delsuc F. 2018. MACSE v2: Toolkit for the Alignment of Coding Sequences Accounting for Frameshifts and Stop Codons. Molecular Biology and Evolution 35: 2582–2584. doi: 10.1093/molbev/msy159

Ross MT et al. 2009. UKPMC Funders Group The DNA sequence of the human X chromosome. Genome 434: 325–337. doi: 10.1038/nature03440.The

Saenko VS, Chouteau M, Prunier FP, Blugeon C, Joron M 2019. Unravelling the genes forming the wing pattern supergene in the polymorphic butterfly *Heliconius numata*. EvoDevo 10: 16. doi: 10.1186/s13227-019-0129-2

Schwander T, Libbrecht R, Keller L. 2014. Supergenes and complex phenotypes. Current Biology. 24: R288–R294. doi: 10.1016/j.cub.2014.01.056

Sharp PM, Averof M, Lloyd AT, Matassi G, Peden JF 1995. DNA sequence evolution: the sounds of silence. Philosophical transactions of the Royal Society of London. Series B, Biological sciences 349: 241–247. doi: 10.1098/rstb.1995.0108

Skaletsky H, Kuroda-Kawaguchl T, Minx PJ, Cordum HS, Hlllier LD, Brown LG, Repplng S, Pyntikova T, All J, Blerl T, Chinwalla A, Delehaunty A, Du H, Fewell G, Fulton L, Fulton R, Graves T, Hou SF, Latrielle P, Leonard S, Mardis E, Maupin R, McPherson J, Miner T, Nash W, Nguyen C, Ozersky P, Pepin K, Rock S, Rohlfing T, Scott K, Schultz B, Strong C, Tin-Wollam A, Yang SP, Waterston RH, Wllson RK, Rozen S, Page DC. 2003. The male-specific region of the human Y chromosome is a mosaic of discrete sequence classes. Nature 423: 825–837. doi: 10.1038/nature01722

Soojin Y, Charlesworth B. 2000. Contrasting patterns of molecular evolution of the genes on the new and old sex chromosomes of *Drosophila miranda*. Molecular Biology and Evolution 17: 703–717.

Steinemann M, Steinemann S 1992. Degenerating Y chromosome of *Drosophila miranda:* A trap for retrotransposons (chromosome structure/larval cuticle protein genes). Genetics 89: 7591–7595.

Stolle E, Pracana R, Howard P, Paris CI, Brown SJ, Castillo-Carrillo C, Rossiter SJ, Wurm Y. 2019. Degenerative expansion of a young supergene. Molecular Biology and Evolution 36: 553–561. doi: 10.1093/molbev/msy236

Sun Y, Corcoran P, Menkis A, Whittle CA, Andersson SGE, Johannesson H. 2012. Large-scale Introgression shapes the evolution of the mating-type chromosomes of the filamentous Ascomycete *Neurospora tetrasperma*. PLOS Genetics 8: e1002820. doi: 10.1371/journal.pgen.1002820

Suzuki H, Brown CJ, Forney LJ, Top EM. 2008. Comparison of correspondence analysis methods for synonymous codon usage in bacteria. DNA Research 15: 357–365. doi: 10.1093/dnares/dsn028

The UniProt C. 2011. Ongoing and future developments at the universal protein resource. Nucleic Acids Research 39: 214–219.

Vicario S, Moriyama EN, Powell JR. 2007. Codon usage in twelve species of Drosophila. BMC Evolutionary Biology 7: 1–17. doi: 10.1186/1471-2148-7-226

Wang J, Wurm Y, Nipitwattanaphon M, Riba-grognuz O, Huang Y-c, Shoemaker D, Keller L. 2013. A Y-like social chromosome causes alternative colony organization in fire ants. Nature 493: 664. doi: 10.1038/nature11832

Weber CC, Boussau B, Romiguier J, Jarvis ED, Ellegren H. 2014. Evidence for GC-biased gene conversion as a driver of between-lineage differences in avian base composition. Genome biology 15: 549. doi: 10.1186/s13059-014-0549-1

Westergaard M 1958. The mechanism of sex determination in dioecious flowering plants. Adv. Genet. 9: 217–281.

Whittle CA, Sun Y, Johannesson H. 2011. Degeneration in codon usage within the region of suppressed recombination in the mating-type chromosomes of *Neurospora tetrasperma*. Eukaryotic Cell 10: 594–603. doi: 10.1128/EC.00284-10

Wilson MA. 2021. The Y chromosome and its impact on health and disease. Human Molecular Genetics: ddab215. doi: 10.1093/hmg/ddab215

Wint R, Salamov A, Grigoriev IV. 2022. Kingdom-wide analysis of fungal transcriptomes and tRNAs reveals conserved patterns of adaptive evolution. Mol Biol. Evol. https://doi.org/10.1093/molbev/msab372

Wu M, Moore RC. 2015. The evolutionary tempo of sex chromosome degradation in *Carica papaya*. Journal of Molecular Evolution 80: 265–277. doi: 10.1007/s00239-015-9680-1

Yan H, Jin W, Nagaki K, Tian S, Ouyang S, Buell CR, Talbert PB, Henikoff S, Jiang J. 2005. Transcription and histone modifications in the recombination-free region spanning a rice centromere. Plant Cell 17: 3227–3238. doi: 10.1105/tpc.105.037945

Yang Z, Nielsen R. 2000. Estimating synonymous and nonsynonymous substitution rates under realistic evolutionary models. Molecular Biology and Evolution 17: 32–43. doi: 10.1093/oxfordjournals.molbev.a026236

Zhou Z, Dang Y, Zhou M, Li L, Yu C-h, Fu J, Chen S, Liu Y. 2016. Codon usage is an important determinant of gene expression levels largely through its effects on transcription. 9.

